# Nanopore direct RNA sequencing (DRS) of MS2 bacteriophages in *E. coli* throughout its life cycles reveals a complex transcriptional activity to control and maintain its growth

**DOI:** 10.1101/2025.03.30.646209

**Authors:** Ni Ni, Gaetan Burgio

## Abstract

The RNA bacteriophage MS2 is an RNA phage that infects the bacterium *E. coli* and is one of the most studied and prototypical model phages in molecular biology and microbiology. Previous research revealed complex translational control and fine-tuning for MS2 replication. However, the dynamics of its transcriptional activity and replication during the life cycles within the bacteria remain elusive. Here, we employed Nanopore Direct RNA sequencing (DRS) to investigate the transcriptome and epitranscriptome landscape of the MS2 in infected *E. coli* throughout multiple life cycles. We discovered that MS2 phages sustain a high level of transcriptional activity required for replication. We found large amounts of subgenomic small transcripts from RNA degradation, Nanopore DRS bias, and transcripts containing the *coat*-encoding region, required for virion assembly. We found hybrid reads due to the error-prone activity of the MS2 replicase complex by the template-switching mechanism. We evidenced that RNA modification is conserved throughout the entire life cycle in full-length transcripts without the acquisition of new modifications, whereas small transcripts did acquire newly modified sites. The conserved sequence and secondary structure (U-rich hairpin) of Ψ installation sites were the most amenable to RNA modification, and one site aligned well with host RluA-mediated installation. Overall, our investigation reveals a more complex transcriptional dynamics of MS2 phages than anticipated within *E. coli* to maintain its growth and replicate under host pressure.

## Introduction

Viruses of bacteria (bacteriophages) are the most abundant organisms on Earth (Clokie et al., 2011). Phages play an important role in the regulation of biological processes and have been harnessed for biotechnology applications (Zhang and Wu, 2020). RNA bacteriophages (commonly known as RNA phages), such as MS2, R17, PP7 and Q-beta, are single-stranded RNA viruses that infect a variety of bacteria (Bollback and Huelsenbeck, 2001). It is thought that the ssRNA phages are the oldest lineage of RNA viruses still in existence (Wolf et al., 2018). These have proven to be useful models for comprehending basic molecular mechanisms such as RNA genome replication, translational control, and gene regulation (Poot et al., 1997; Rolfsson et al., 2016).

MS2 belongs to the family *Fiersviridae*, and is an icosahedral, non-enveloped, F-specific, positive-sense single-stranded RNA (+ssRNA) lytic coliphage (Salas & de Vega, 2008). The ssRNA phages adopt a simple infectious cycle that includes host adsorption, genome entry (genomic RNA penetration), genome replication, phage assembly, and host lysis to release new bacteriophage progeny for successive infections. MS2 phage replicase β subunit and host proteins (ribosomal protein S1 subunit, EF-Tu subunit (elongation factor thermo unstable), and EF-Ts subunit (elongation factor thermos stable)) form an RNA-dependent RNA polymerase (Rdrp) complex required for replication (**Figure 1A**) (Wagner, Weise, and Mutschler, 2022). Full-length negative single-stranded RNA intermediates are generated to serve as the template for the synthesis of positive genomic RNA (gRNA) by the replicase complex (**Figure 1A**). Among phages, ssRNA phage MS2 has one of the smallest and simplest genomes with only possessing four ORFs in its 3,569 nt genome that also serves as an mRNA: *mat* encodes maturation protein (MP), which enables attachment to the F pilus and RNA entry into the host (Meng et al., 2019); *coat* encodes the coat protein (CP), which forms the viral capsid; *lys* encodes lysis protein (L), which allows for host lysis (Chamakura, Tran, and Young, 2017); and *rep* encodes catalytic subunit of replicase (Rep), which is an RNA-dependent RNA polymerase (Wagner, Weise, and Mutschler, 2022) **(Figure 1B)**.

**Figure 1.**
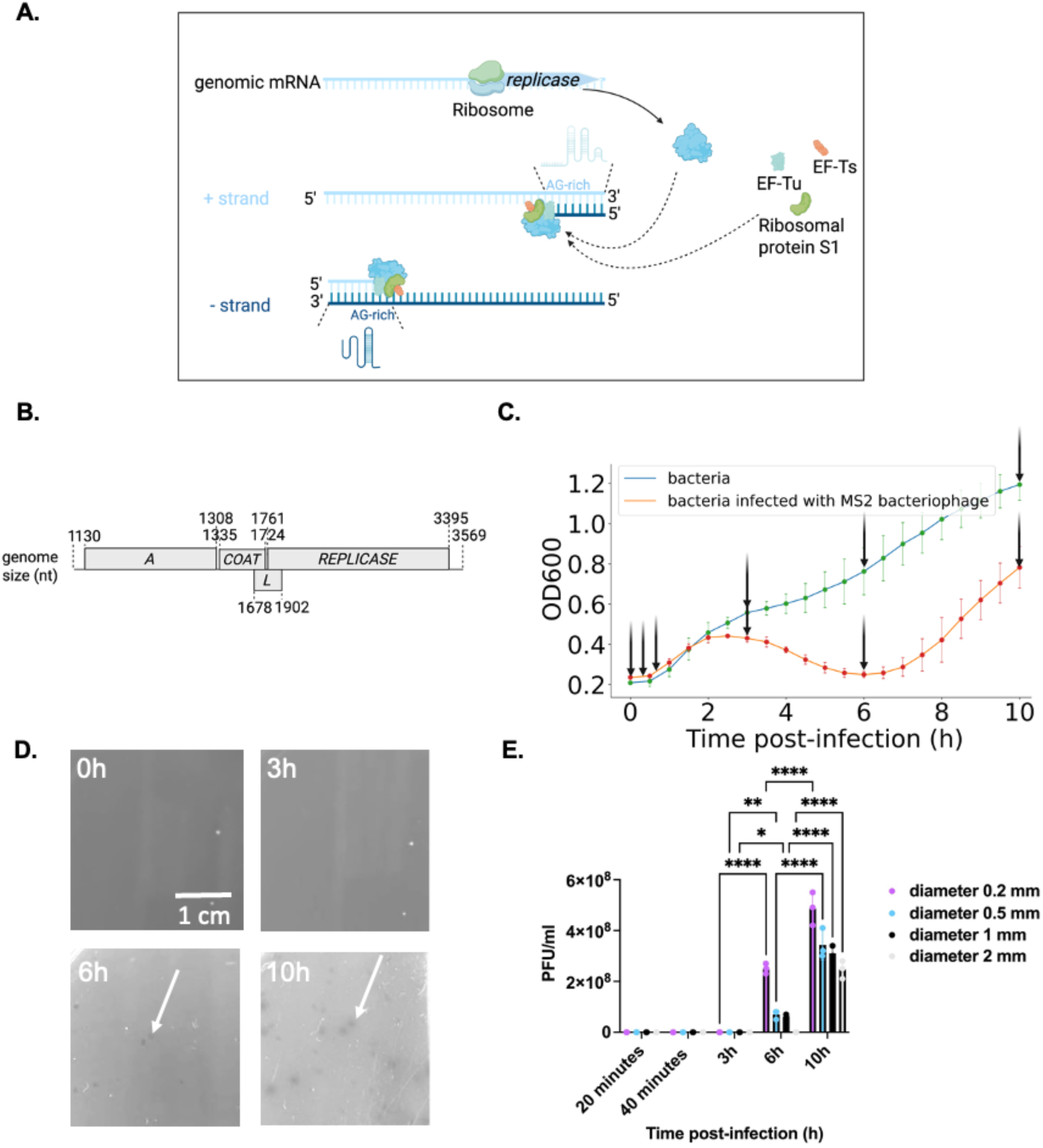
Characteristics of MS2 phage. **A.** MS2 replication pattern. The MS2 replicase complex contains host factors EF-Tu, EF-Ts, and S1 ribosomal protein. **B.** Schematic presentation of the MS2 full-length genomic RNA coding regions organisation and composition. *A*, maturation protein-encoding gene; *COAT*, coat protein-encoding gene; *L*, lysis protein-encoding gene; *REPLICASE*, replicase protein-encoding gene. NTD, N-terminal domain; CTD, C-terminal domain. **C.** Growth curves of bacterial cells (DSM5695 *E. coli*) infected with MS2 RNA phage at MOI = 200 (red line) and bacterial cells without infection (blue). **D.** MS2 plaques on the host *E. coli* DSM5695. Plaques were pointed with white arrows. **E.** Plaque PFU and size change. All pair-wise comparisons were performed after observing a significant Analysis of Variance (ANOVA) with * *p* < 0.05, ** *p* < 0.01, **** *p* < 0.0001. Data are presented as means ± SD (n = 3).

Previous research revealed RNA–protein interactions that regulate the precise timing and strength of viral protein production throughout the bacteriophage life cycle (Dykeman, 2024; Twarock et al., 2024). Despite years of research on MS2 replication and assembly, little is known about the dynamics of transcription of MS2 during its life cycle. Like many other viruses, MS2 RNA encodes an error-prone replicase (Wagner, Weise, and Mutschler, 2022). The replicase complex is repressed from the CP, which binds to specific sites on the positive-strand RNA (Peabody, 1990). While the interplay between different phage proteins resulting in entry, replication, assembly, and encapsidation has been relatively well studied in *in vitro* settings (Rolfsson et al., 2016; Wagner, Weise, and Mutschler, 2022), the dynamics in transcription throughout the MS2 life cycle remain elusive. Specifically, the unanswered questions remain on how MS2 transcriptional activity rapidly increases to sustain the production of the virion, the nature and the extent of the byproducts the error-prone replicase produces during the MS2 life cycle, and how the dynamics of acquisition of *de novo* mutations and RNA modification occur in the bacterial cell.

To investigate the MS2 transcriptome dynamics across time, Direct RNA sequencing using Oxford Nanopore sequencing enables the sequencing of long RNA sequences encompassing the full-length transcript and the identification of RNA modification in infected cells. Previous work has provided valuable insights into the dynamics of transcription, RNA splicing (Kim et al., 2020), recombination (Viehweger et al., 2019), and RNA modification landscapes of various viruses (Viehweger et al., 2019).

Here, we employed Nanopore DRS to investigate the transcriptome and epitranscriptome landscape of the MS2 in infected *E. coli* throughout its life cycle. We discovered that MS2 phages sustain a high level of transcriptional activity within the bacterial cell. We found sub-genomic transcripts resulting from either RNA degradation or Nanopore DRS bias. Interestingly, we found the presence of small sub-genomic transcripts encoding the full-length coat protein ORF. The MS2 replicase is error-prone, which frequently generates hybrid RNAs across time by the template-switching mechanism. We evidenced that RNA modification is conserved throughout the entire life cycle in full-length transcripts without the acquisition of new modifications, whereas small transcripts did acquire modified sites. The conserved sequence and secondary structure (U-rich stem-loops) of Ψ installation sites were the most amenable to RNA modification and likely mediated by host pseudouridine synthase RluA. Together, using DRS RNA Nanopore sequencing, we have dissected the dynamics of MS2 transcriptional activity in bacterial cells across time.

## Methods

### Bacteria, phage strains, and culture conditions

The bacteria host *E. coli* DSM 5695 (also called *E. coli* W1485) (F^+^) and MS2 bacteriophages DSM 13767 were obtained from the DSMZ-German Collection of Microorganisms and Cell Culture. The bacteria were maintained in NZCYM media (10.0 g/L Casein hydrolysate, 5.0 g/L NaCl, 1.0 g/L Casamino acids, 5.0g/L Yeast extract, 2.0 g/L MgSO4 X 7H2O) with maltose (2g/L) and incubated at 37°C overnight. MS2 phages were maintained and propagated in SM buffer (NaCl 100 mM, MgSO4 X 7H2O 8 mM, Tris-HCl 50 mM, gelatin 0.01% (w/v), pH 7.5). MS2 plaque assays were performed in a double-layer Agar. Briefly, 100 µl overnight culture of DSM 5695 bacteria and 100 µl MS2 in SM buffer were mixed with 5 ml NZCYM soft agar with maltose (2g/L). Plates were incubated at 37°C overnight (∼16 h). MS2 phages were harvested. The top layer was disrupted after pouring 5 ml SM buffer, and the lysate was centrifuged at 2,800g for 30 minutes at room temperature. The lysate was filtered using a 3 ml syringe with a 0.22 µM filter. The precise viral stock titer (PFU/ml) of the phage (MS2) was determined.

### MS2 phage infection and RNA isolation

The bacteria *E. coli* (DSM 5695) liquid culture growth infected with bacteriophage MS2 was generated by multimode plate reader IID VICTOR Nivo (PerkinElmer), start OD 600 of overnight DSM 5695 bacteria culture was around 0.2-0.3. Each data point was the result of at least three biological replicates. DSM 5695 *E. coli* cells (overnight culture) were infected with MS2 at a multiplicity of infection (MOI) of 200 and incubated for 6 h at 37 °C. Infected DSM 5695 cells and non-infected DSM 5695 cells (i.e., mock-infected cells) were harvested (∼10^10^ bacterial cells were prepared as the initial material) at 0h, 20 and 40 minutes, 3h, and 6h post-infection (p.i.). At the 0h time point, immediately after adding the phages, we isolated the total RNA, and the bacterial cells were not washed. Total RNAs were extracted from infected *E. coli* samples (TRIzol [Invitrogen]) following the manufacturer’s instructions.

### Reverse transcription quantitative PCR (RT-qPCR) of MS2 phage genes

The primer pairs for RT-qPCR were designed specifically to the MS2 gene regions using RefSeq Representative Genome Database (Organism limited to *Escherichia coli*) with the NCBI Primer-BLAST tool. The *E. coli cysG* gene (housekeeping gene) was selected as the reference gene. Briefly, reverse transcription for 100 ng whole RNA extracted was performed with SuperScript™ IV Reverse Transcriptase [Invitrogen] with random primers (hexamers) [Invitrogen] at 50°C, then qPCR was performed with PowerTrack™ SYBR Green [Invitrogen] using specific sets of primers (**Suppl. Table S12**) for specific phage genes (400 nM primers with annealing temperature 60°C) with input cDNA (serially diluted to determine the qPCR primer efficiency) according to the manufacturer’s protocol an visualized by QuantStudio™ 12K Flex Real–Time PCR System. Data analysis using the ΔΔCT method was performed to quantify the relative expression of MS2 phage genes.

### *In vitro* transcription of MS2 RNA

One μg of total RNA was purified from MS2-infected *E. coli* cells. Reverse transcription (SuperScript™ IV Reverse Transcriptase [Invitrogen]) was performed with each phage-specific RT primer (RTprimer1-6) (**Suppl. Table S12**). Templates for *in vitro* transcription were prepared by PCR with each phage-specific PCR primer pair (i.e., IVT-frag1-F & IVT-frag1-R primers with 5’ terminal T7 RNA polymerase promoter region) followed by agarose gel purification (Wizard SV Gel and PCR Clean-Up System kit [Promega]), *in vitro* transcription using T7 polymerase (MEGAshortscript™ T7 Transcription Kit [Thermo Fisher]) and RNA purification (Monarch RNA Cleanup Kit [NEW ENGLAND Biolabs]) with re-suspension in ddH2O. The oligonucleotides used in this study are listed in the **Suppl. Table S12**.

### Poly(A) tailing of RNA and beads purification

Total RNAs and IVT RNAs were heat incubated at 65°C for three min and cooled on ice for one min before polyadenylating RNAs using the *E. coli* poly(A) polymerase (NEB M0276S [NEW ENGLAND Biolabs]). Briefly, 10 µg RNA, 5 units poly(A) polymerase, 2 µL 10X reaction buffer, and 1 mM ATP in a total reaction volume of 20 µL were incubated for 45 min at 37°C. To stop and clean up the reaction, poly(A)-tailed RNAs were purified following the Agencourt AMPure XP magnetic beads (1:1.8 RNA to beads ratio) according to the manufacturer’s protocol and resuspended in 10 µL of ddH2O. The integrity of total RNA from *E. coli* was assessed via a Bioanalyzer (Agilent).

### Direct RNA nanopore sequencing

The Direct RNA Sequencing Kit (SQK-RNA004 by Oxford Nanopore) was used to prepare the libraries and sequence the native poly(A)-tailed RNA library according to the manufacturer’s protocol and including the reverse transcription step to generate RNA:cDNA hybrids. Sequencing runs were performed using PromethION flow cells (FLO-PRO004RA). The Flow Cell Wash Kit (EXP-WSH004) was used to remove the library between flow cell washes and library reloading, following the manufacturer’s instructions.

### Basecalling and read mapping

The Computational workflow for processing the Oxford RNA-direct Nanopore sequencing datasets was shown (**Suppl. Figure S1**). Basecalling was performed using MinKNOW software (Nanopore) using the Super accurate model. The Fast5 files were filtered. Reads with a minimum PHRED score of 10 were removed. Fast5 files were then converted to pod5 using pod5_file_convert script using the –one-to-one option. Basecalling on pod5 files was performed with Dorado (v 0.9.1) using the rna004_130bps_sup@v5.1.0 basecalling model for m5C, m6A_DRACH, Inosine_m6A, PseuU. Reads were mapped to the reference sequence NC_001417.2 using minimap2 (Li, 2018) using the following parameters: -ax map-ont -y --secondary=no -k 4 -w 10. Samtools (Li et al., 2009) was used to retain the mapped reads either on the positive strand (Samtools view –F2324 -b) or negative strand (Samtools view –f 16 –b) and reverse complemented. Reads were sorted and indexed. Mapped read lengths were counted using genomecov function from Bedtools (Quinlan and Hall, 2010). Other metrics, such as alignment identity, were determined using MarginStats (Jain et al., 2015) and based on a similar workflow previously described (Workman et al., 2019).

### Gene expression analysis

To map the reads to transcripts, a transcript reference sequence was generated using the fasta reference sequence and a gff file converted from genebank format. Reads were mapped to the transcript using minimap2 with the same options as previously. Mapped reads were counted with Salmon (Patro et al., 2017) using the options: --ont –P 48 –l U. Gene expression analysis was performed with Deseq2 (Love, Huber, and Anders, 2014) under R 4.4.0 using time as a variable. Gene expression was normalised and compared for each gene across time.

### Read coverage analysis

Read length was determined with Samtools and separated into multiple fractions of similar read lengths. Reads were mapped to the NC_00141702.2 genome (DSM 13767 genome assembly lacked the 3’ end of the contig, preventing its use as a reference genome) and sorted using Samtools. Reads were also mapped to the bacterial host strain DSM 5695 for read count. Read coverage within a 5 bp interval for reads starting/ending versus the total number of read at these specific window positions using Samtools and a custom bash script. Stop and start codons were determined using a motif search and overlayed to the relative coverage plot. The secondary RNA structure was determined using RNAfold. Hybrid reads were mapped to the NC_00141702.2 genome using blastn. The top hits e.values > 10^8^ were retained. The query adjacent reads were retained for further analysis, and the target read locations were determined on the positive and negative strands and mapped back to the NC_00141702.2 genome.

### Detection of single nucleotide and structural variants

To detect single nucleotide variants and indels, we performed the analysis using Clair3 (Zheng et al., 2022) and Sniffles2 (Smolka et al., 2024). We included the following flags for Clair3 analysis: --platform=”ont” --model_path=r941_prom_sup_g5014 –include_all_ctgs – no_phasing_for_fa. For indels detection, we performed the analysis using Sniffles2 with default parameters. The files were filtered and converted using Bcftools (Danecek et al., 2021) in vcf format.

### RNA modification analysis

Basecalled reads from Dorado were mapped to the NC_00141702.2 genome using Minimap2, sorted, and indexed using Samtools. Basecalling for modification was determined using Modkit pileup function using –max-depth 2000000 option and a modification threshold > 80%. Differential methylation profile was determined using modkit dmr function. The Nanopore DRS technique has the limitation of high false-positive rates (FPR), and the IVT samples were prepared to reduce the FPR. Only Stoichiometry (test) - Stoichiometry (IVT) > 0 was included in our study.

### Statistical analysis

Statistical analysis was conducted. The mean ± standard deviation (SD) was shown. The differentially expressed genes were identified by assessing the statistical significance via the adjusted *p*-value after accounting for multiple hypothesis testing (Benjamini-Hochberg method). The threshold for statistical significance was set at *P* value < 0.05, *, *P* value <0.01, **. The Benjamini-Hochberg method was performed to control the false discovery rate (FDR correction) for post-transcriptional modifications (Benjamini and Hochberg, 1995). Graphs were generated, and data were visualized using ggplot2 from R (version 4.4.0), GraphPad Prism (version 10.4.1), and Matplotlib from Python (version 3.12.8).

### Sequencing data Files and code availability

Raw sequencing data files are available at the NCBI Sequence Read Archive (SRA) under the following accession ID: Bioproject ID PRJNA1243726 and BioSamples SAMN47628658 to SMNA47628577. The custom source codes used for the computational analysis have been deposited in the following Github repository: https://github.com/gburgio/MS2_Nanopore_DRS.

## Results

### MS2 bacteriophage replicated within half an hour in *E. coli*

To determine the course of infection in *E. coli* host over 10 hours post inoculation (p.i.), we infected the bacterial cells with MS2 phage and monitored *E. coli* growth over time in liquid culture at the multiplicity of infection (m.o.i) of 200 (**Figure 1C**). MS2 formed plaques from 6 hours p.i., indicating its lytic nature (**Figure 1D**). The size of MS2 plaques ranged from 0–2 mm in diameter within 10 hours (**Figure 1 D-E**). Overall, the data indicated that the time points 20 minutes (before the first burst), 40 minutes (after the first burst), 3h (maximum growth), and 6h (lowest growth) p.i. captured a single and multiple life cycles, resulting in plaque formation. We chose these time-points to assess the dynamics of phage transcription across a single and multiple life cycles.

### MS2 phage maintained a high transcriptional activity across time

To capture MS2 transcriptional changes across time, we performed long-read Nanopore direct RNA sequencing (DRS) runs on total RNA extracted from DSM5695 *E. coli* cells infected with MS2. We mapped over 17,6 million reads over a PHRED quality score threshold of 10. Out of 7.5 million reads were mapped on the MS2 genome, we found an average read length spanning from 668 to 954 nt (**Suppl. Table S1)**. Reads over 90% of the MS2 transcript were included as full-length reads (**Suppl. Table S1**). Immediately after infection (0h p.i.), only full-length RNAs were detected (which accounted for only 1.63% of the total viral reads) (**Figure 2A and Suppl. Table S2**) and almost no observable negative-strand transcripts (**Suppl. Figure S2C)**. A sudden increase in the ratio of viral RNAs to bacterial RNAs occurred from 20 minutes (1.05%) to 40 minutes (56.13%), suggesting the first cycle burst occurred at ∼30 to 40 minutes post-inoculation (**Figure 2A and Suppl. Table S2)**. Gene expression analysis was conducted to assess variation between different time points. From the Principal Component Analysis (PCA), the observed variation on the first two PC axes accounted for 88% of the total variance (**Figure 2B**). Three main groups were identified: the 0h and 20 minutes p.i. on the PC1 negative axis and the infected group over 40 minutes p.i. on the positive PC1 axis (**Figure 2B**). Among the infected group, the 40-minute time point group was on the PC2 positive axis, whereas the 3 and 6 hours p.i. were clustered on the negative PC2 axis **(Figure 2B)**. The different time points exhibited distinct clusters on the correlation heatmap (**Suppl. Figure S2A**), confirming significant transcriptional changes of MS2 across time. Next, we analyzed the differential expression (DE) of the four MS2 ORFs across time (**Figure 2C**). We found an upregulation of the *mat* and *cap* ORFs while the *rep* ORF was downregulated (**Figure 2C**). Interestingly, we found a peak expression of the *lys* gene at 40 minutes p.i., capturing a potential phage burst. We confirmed by RT-qPCR similar results to the Nanopore DRS (**Suppl. Table S3**). Compared with 0h p.i., we importantly found a significant fold increase of > 400-fold at 3 and 6 hrs p.i. of the coat protein (**Suppl. Table S3**). The results also indicated the relatively higher expression of phage RNA of the coat-protein ORF than the maturation-protein ORF at the same time point (**Suppl. Table S3**), confirming our observations from Nanopore DRS. Overall, it suggests a significant up regulation of the coat protein and fine tuning of the ORFs expression to regulate MS2 transcription across time.

**Figure 2.**
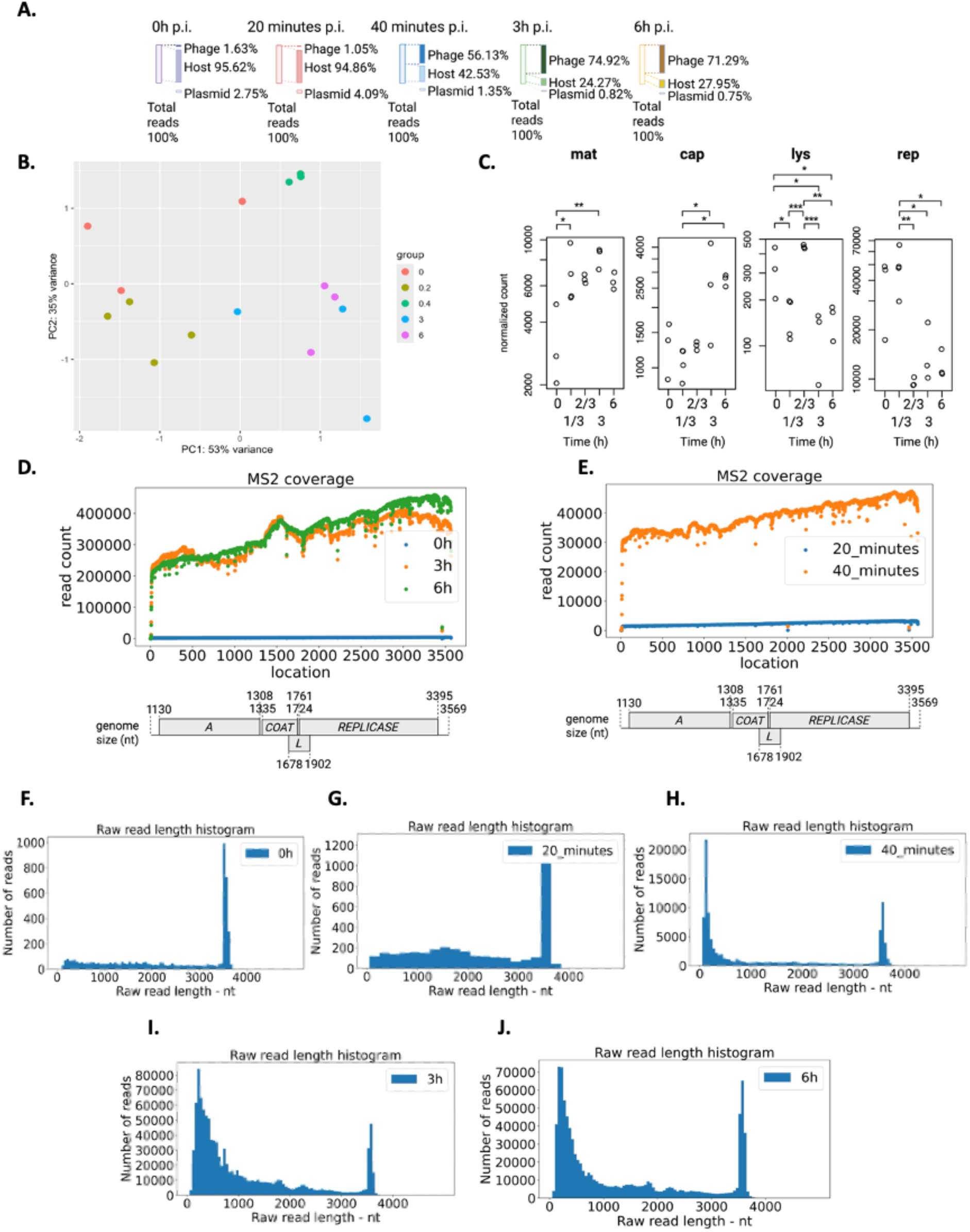
Statistics of Oxford Nanopore Technologies (ONT) direct RNA sequencing data via computational approaches and transcription pattern of MS2. **A.** Read counts from Nanopore direct RNA sequencing of total RNA from DSM5695 *E. coli* cells infected with MS2 phage at different time points. **B**. Principal component analysis (PCA) plot of each MS2 sample. **C**. Differential expression patterns for four MS2 genes at different time points based on normalised read count. * *p* <0.05, ** *p* < 0.01, *** *p* < 0.001. **D**. Coverage map for MS2 total reads at 0 to 6 hours p.i., **E**. 20 and 40 minutes p.i., **F**. MS2 raw read length distribution at 0h p.i. **G**. At 20 minutes p.i. **H**. At 40 minutes p.i. **I**. At 3h p.i. J. At 6h p.i.

### Read mapping revealed a bias in coverage towards the coat protein ORF

During replication, MS2 forms an intermediate negative transcript that serves as a template for transcription/translation of the ORFs (Wagner, Weise, and Mutschler, 2022). We separated the positive and negative strands, mapped reads to capture the intermediate state of viral replication. Next, we assessed the read coverage across time for all reads (**Figure 2D-E**), positive and negative strands (**Figure 3A-B**). We observed a coverage bias towards the *coat* ORF at 3h and 6h p.i. and the 3’ end of the viral genome on the overall transcript (**Figure 2D),** further confirming our results. As the nanopore DRS has the 3’ bias due to directional sequencing from the 3’ ends of RNAs (Jain et al., 2022), we deduced that coverage was biased only towards the *coat* ORF in the MS2 transcriptome.

**Figure 3.**
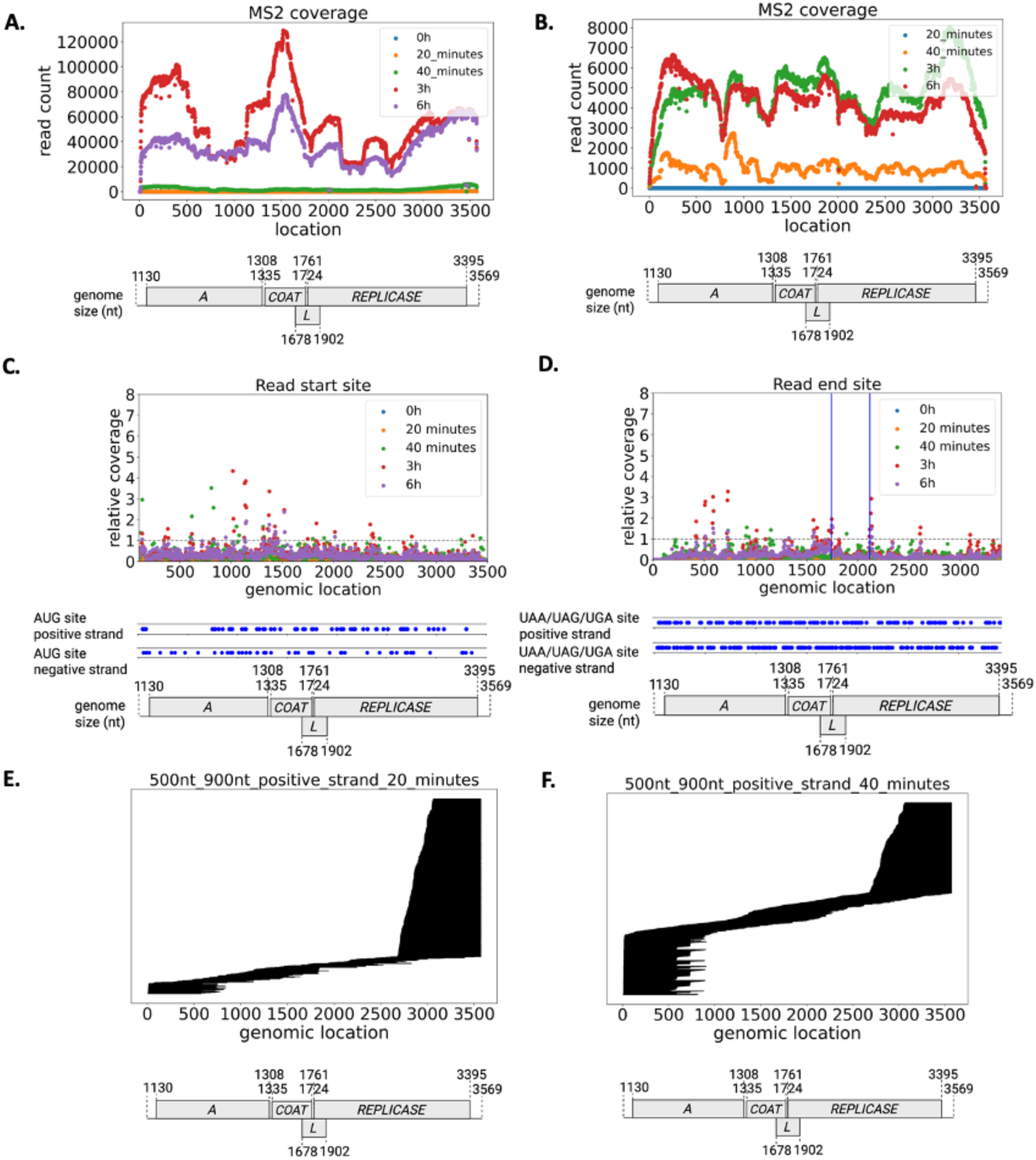
Replication pattern of MS2. **A.** Coverage map for MS2 reads (< 800 nt) aligned to the positive strand and **B.** the negative strand. **C.** Relative read start site (ranging 100-3500 nt) coverage profile and **D.** Relative read end site (ranging 0-3400 nt) coverage profile for MS2. The blue vertical line indicated two previously identified key coat protein binding sites. **E**. Positive-strand reads (shown as lines) aligned to the MS2 genome with read length between 500 nt and 900 nt at 20 minutes p.i., and **F.** Read length between 500 nt and 900 nt at 40 minutes p.i.

### Emergence and dominance of short transcripts across time

While quantifying read counts, we noticed over 60-fold increase of the full-length transcripts (> 3000 nt) across time (**Figure 2 F-J**), consistent with the gene expression data. Intriguingly, we noticed the emergence and dominance of a large population of short reads (< 800 nt) from 40 minutes p.i. (**Figure 2 H-J and Suppl. Figure S2 F-K)** on positive strand, ranging from 37 % to 55 % of the read count (**Suppl. Table S4**). A possible explanation is that these reads are the accumulation of degraded RNA via bacterial lysis. As such, the number of these short reads would increase in the phage and the bacteria across time. We quantified the proportion of these short reads in the host genome, and we found a similar percentage variable from 82% to 92% of the total read count, with a decrease to 82 - 85% from 20 minutes to 3 hours p.i. (**Suppl Figure S4 and Suppl Table S5**). It suggests that the increase in the proportion of the short-read count found in phage is not correlated with bacterial lysis. To quantify a potential RNA degradation of these reads mapped to the phage transcript, we performed a quality control of the samples and the reads consisting in measuring the RNA integrity number (RIN) (**Suppl. Table S6**), the quality score versus read length and the number of mismatches per read to the reference transcript, proxy for read quality (**Suppl Figure S3**). The RIN score shows a low amount of RNA degradation with a median of 7.8 ±0.6 (**Suppl. Table S6)**. To miminise the effect of RNA degradation in our analysis, we opted for a more stringent cutoff from the literature and retained reads over a PHRED score of 10. While N50 statistics show a slight reduction in the N50 length across time, from a median length dropped from 1,432 nt at 0 hrs p.i. to 1,185 nt at 6 hrs p.i.

### AG-rich regions were responsible for the initiation of small transcripts

Previous works described the ability of the replicase to generate small spontaneous template-directed RNA species (Chetverin, Chetverina, and Munishkin, 1991; Wagner, Weise, and Mutschler, 2022). We hypothesized that these small transcripts could be either RNA species resulting from Nanopore DRS 3’ bias and library preparation, small transcripts required for replication, and/or products of RNA degradation from bacterial lysis.

To further investigate the nature of these small transcripts, we quantified their coverage across the MS2 genome (**Figure 3 A-B**). We found an enrichment over 2-fold of the 5’ end and the *coat* ORF (∼18.8% of total positive-strand reads with read length <800 nt at 3h) on the positive strands (**Figure 3A and Suppl. Table S7),** confirming the initial mapping and RT-qPCR results (**Suppl. Table S3**). We also found enrichment of the 3’ end of the full transcript, likely due to the DRS bias. Interestingly, transcripts mapped to the 3’ end of the *rep* ORFs were enriched on the negative strand (**Figure 3B**), whereas part and the 3’ end of the *mat* ORF lacked coverage on the positive strand (**Figure 3A**). To detail the mapping of these < 800 nt transcripts, we performed a genome-wide mapping of the start and end of these transcripts within a 5 bp size window and calculated the fold changes relative to the overall coverage. We noticed peaks of significance on the positive and negative strands (**Figure 3 C-D**). We notably observed hotspots at ∼1,000-1,300 bp within the *mat* ORF and hotspots of end sites at ∼1740-1745 nt after the c*oat* ORF and at 2125-2130 nt within *rep* ORF (**Figure 3 C-D**).

We next postulated that specific RNA signals (ribosome binding sites/codon start) are the key determinants for MS2 transcript initiation of these < 800 nt transcripts. We investigated several start site peaks by assessing the genomic composition and secondary structure of their corresponding genomic locations. We found AG-rich regions near these hotspots on the negative strand. Using Clustal alignment and MEME, we found that the MEME motif (**Suppl. Figure S5 A-E**) of these regions superimposed to known ribosomal binding sites (RBS) (Ng, Wenfa (2019) (**Suppl. Table S8**). Together, they suggest the production of small transcripts during the phage life cycles.

### A population of subgenomic reads is produced during the MS2 life cycles

To replicate its own transcript, MS2 generates a negative-strand transcript, which initiates the replication from the 3’ to the 5’ end (Wagner, Weise, and Mutschler, 2022). Prior work on Nanopore sequencing technology has demonstrated that degraded RNAs/random cleavages during library preparation exhibit a trimmed 3’ end (Prawer et al., 2023). We postulated that transcripts mapped starting from the 3’ end resulted from Nanopore DRS 3’ end bias since the sequence starts from the 3’ end, whereas transcripts mapped from the 5’ end were degraded transcripts. We aligned these short reads to the MS2 transcript. We found three types of aligned reads (**Figure 3E-F**). The first block aligned from 3’ to 5’ end accounted for 78.42% of the read count (length ranging 500nt – 900 nt) at 20 minutes p.i. (**Suppl. Table S9**) and reduced in proportion after the first cycle. These reads corresponded to Nanopore 3’ end bias. The second block aligned from 5’ to 3’ end accounted for degraded RNA and increased over time, likely due to active lysis in bacteria and potentially DRS sample preparation. Previous studies identified that the coat protein subunit binds alongside the RNA template to mediate and regulate the viral assembly and packaging during replication (Valegård et al., 1994). Interestingly, a third block of aligned transcripts aligned to the *coat* and the *lys* ORFs and increased across time (**Figure 3F and Suppl. Figure S6B**). This observation was in accordance with our previous results on the gene expression analysis backed by RT-qPCR (**Figure 2D**). Within one life cycle, we observed an increased count of transcripts containing the coat protein encoding region, from 1.92% for 0h, 2.73% for 20 mins, and 3.38% for 40 mins (**Suppl. Table S9**). For transcripts containing both the coat and lysis protein encoding region, we observed an increase of 0.14% at 20 minutes to 1.23% at 40 minutes (**Suppl. Table S9**). We found no clear alignment patterns with MS2 on the negative strand, suggesting that, unlike the full-length transcripts, these small transcripts did not produce a negative strand (**Suppl. Figure S6 D-F, Suppl. Table S10**). Overall, it suggests that these small transcripts throughout the life cycle(s) vizualised are a combination of degraded RNA, small transcripts required for phage replication, and Nanopore DRS 3’ bias across time.

### Error-prone replicase frequently generates hybrid RNAs

Previous works have described the generation of by-products from the error-prone replicase (Kopsidas et al., 2007). While it was previously reported that MS2 phage was capable of forming hybrid RNAs, it was not found in an *in vivo* setting (van Meerten et al., 2002). Nanopore DRS enables the characterization of recombination events and template switching in viral RNA (Johnson et al., 2024). We interrogated the landscape of hybrid RNA transcripts from DRS. We quantified the presence of hybrid reads by blasting these reads to MS2 genomes and retaining the reads mapping to at least two different MS2 locations in single reads with evidence of a recombination event (**Figure 4A**). Reads of recombination (20-60 nt deletions and/or duplications) within the MS2 RNAs were detected (**Suppl. Table S11**). We found a 0.2% of duplicated or deletion hybrid reads to total reads ratio on the positive strand at 6h p.i. We found a few RNA joining events of the negative strand to negative strand and sporadic negative strand to positive strand, and very rare (+-) foldback RNAs (also known as copyback or snapback RNAs - **Suppl. Figure S7 A-B**). We noticed internal tandem duplication events and deletion events between the negative strand and the negative strand (**Suppl. Figure S7 A)**. While we discerned recombination events within phage RNA, we did not observe any recombination events between viral genome segments and host RNAs. We wondered whether new variants have emerged over time. We assessed the mutational landscape of MS2 phage (SNV and structural variation) across time. We have not observed the emergence of any new mutations up to 6 hours p.i. Together, it indicates the error-prone replicase generates hybrid transcripts and foldback RNAs as by-products.

**Fig 4.**
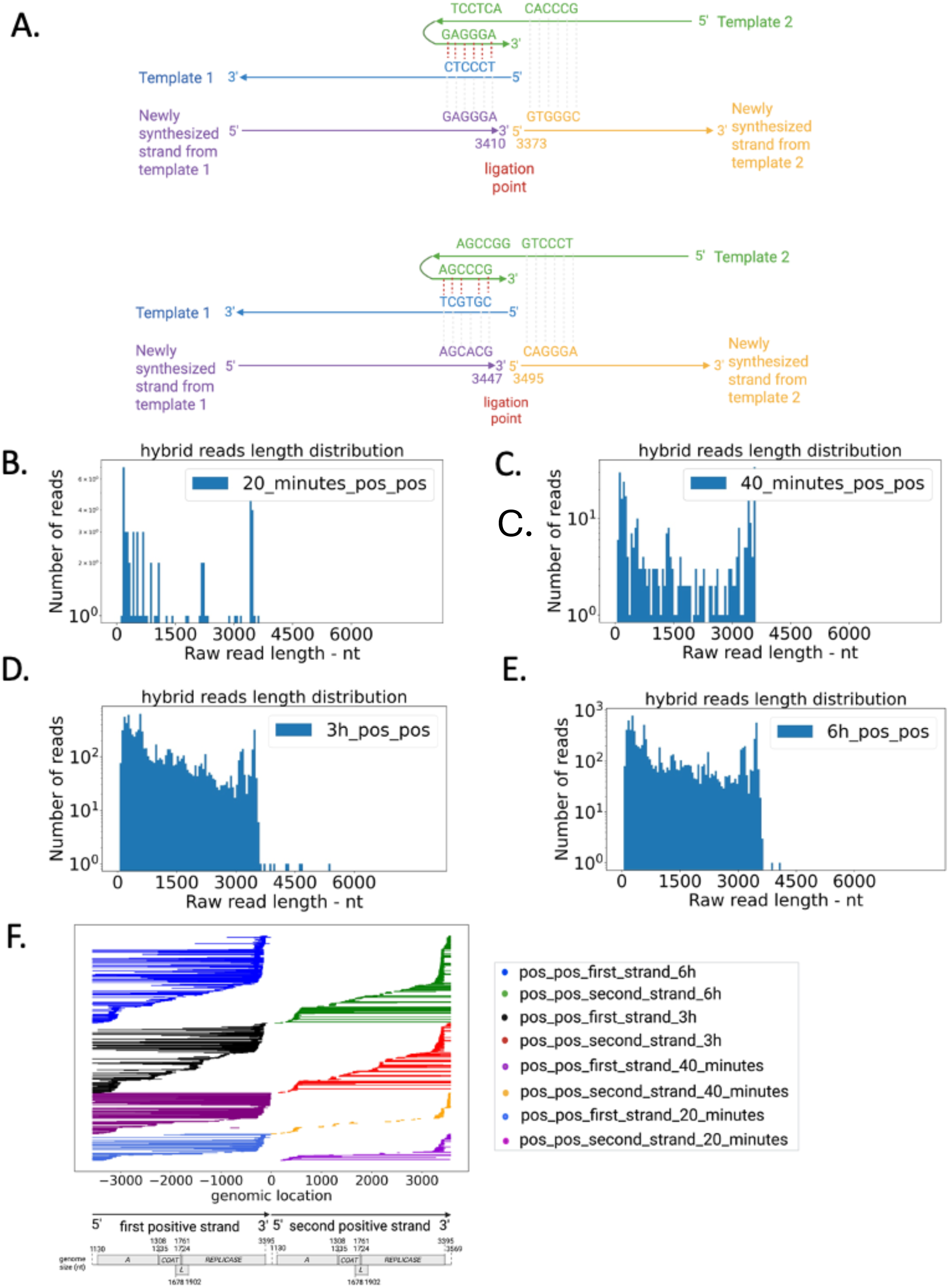
Characteristics summary of MS2 hybrid reads. **A**. Examples of template switching events occurring at multiple time points, ligation at location 3410 and 3373, and ligation at location 3447 and 3495. Template 1 and Template 2 have a complementary region (red dashed line). Replicase first uses template 1 as the template, then switches to template 2 as the template for the following RNA synthesis. **B.** Read length of hybrid reads (positive to positive strand ligation - pos_pos) histogram at 20 mins, **C.** 40 mins, **D.** 3h, and **E.** 6h post inoculation. **F.** Features of hybrid reads (pos_pos) of MS2. Ligation events occurred at the 3’ end of the first strand and the 5’ end of the second strand. All pos_pos hybrid (20 minutes) reads were visualized. 30% of total pos_pos hybrid (40 minutes) reads were visualized. 2% of total pos_pos hybrid (3h and 6h) reads were visualized.

### Static and dynamic RNA modification on MS2 phage during replication

RNA modification plays an important role in modulating RNA function (Pozhydaieva et al., 2024). Previous works have identified the presence of Ψ modification on MS2 phages using either mass spectrometry and/or by DRS (Jones, 2023). We wondered whether MS2 acquired new modifications during a single life cycle (from 0h to 40 minutes p.i.) and across time (3 and 6 hours p.i.). We predicted and mapped m5C and Ψ modifications throughout a single MS2 life cycle on the RNA positive strands (**Figure 5 A-D**). We found m^5^C, and Ψ modifications were widely distributed (**Figure 5 A-D**). On full-length transcripts (> 3,000 nt), we found no evidence of newly acquired modifications within a single cycle (**Figure 5 A and C**). We identified instead commonly modified static sites with high stoichiometry throughout the transcripts, varying from 23.4% to 78.0% (**Figure 5 A and C**). Interestingly, we found shorter transcripts increase in m5C stoichiometry across time (**Figure 5B and Suppl. Figure S9B**). Consistent to a previous study (Jones, 2023), we also identified a conserved Ψ modification site 924 nt (**Figure 5C**) We found additional m5C (at location 634, 927, 2636, 3378, 3413, and 3521 nt) and Ψ installation sites (at location 2145 and 2407 nt) for small transcripts with length < 800 nt at 3h and 6h p.i. (**Suppl. Figure S9 B-D**). These additional modifications were located on the *mat* and *rep* ORFs. We assessed m6A modification, and we observed no modification pattern (**Suppl. Figure S8 A-D**). Overall, it suggests that RNA modifications in modified sites vary in stoichiometry during a single life cycle. Additionally, we demonstrated that *de novo* and stable modifications were acquired across time.

**Figure 5.**
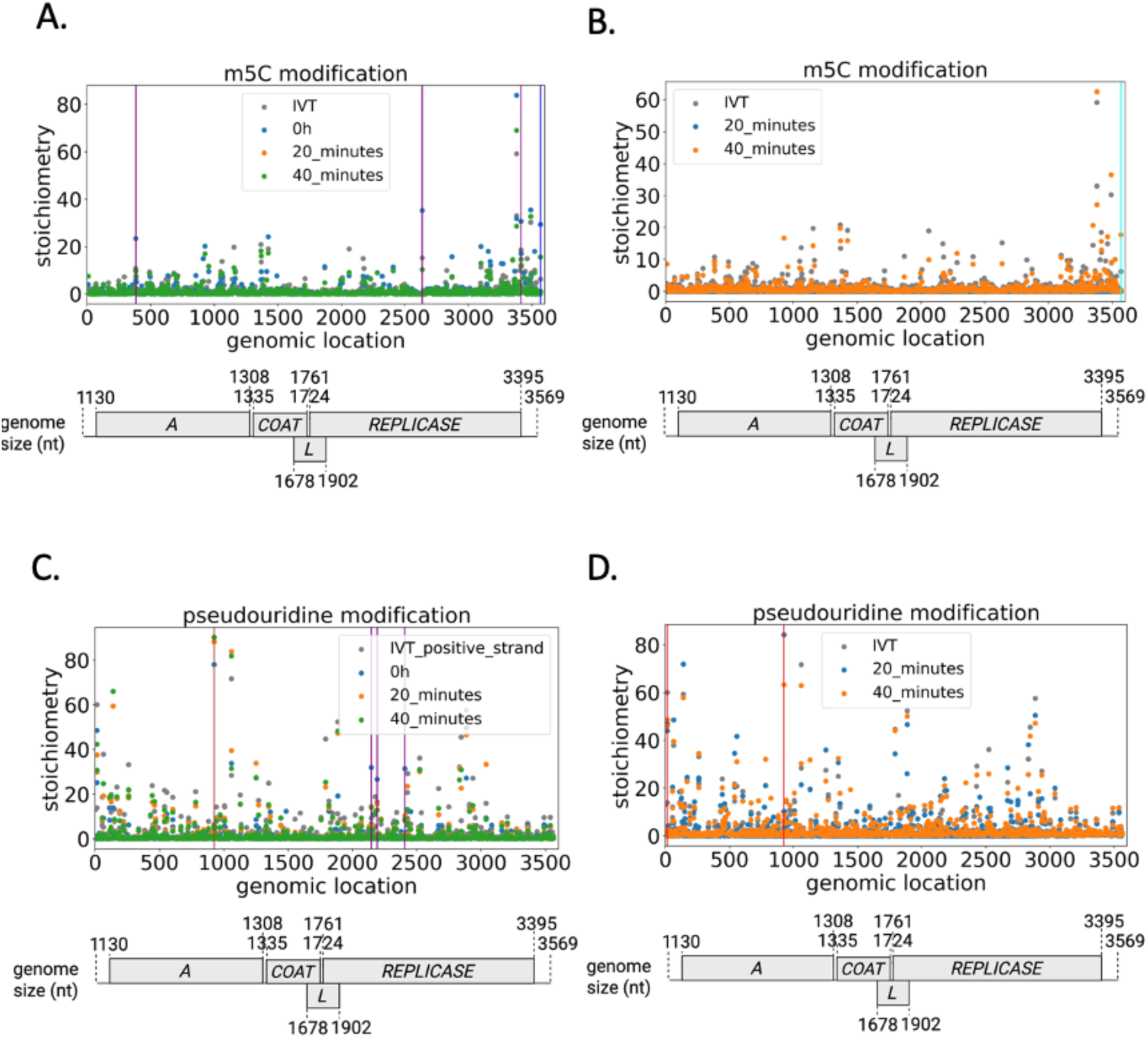
Predicted sites of Ψ and m^5^C RNA modification dynamics on MS2 phage positive RNA (>3000 nt length and <800 nt length) within one life cycle (at 0h, 20 minutes, and 40 minutes p.i.). Synthetic modification-free RNA as the negative control. **A.** m5C modification (>3000 nt) landscapes according to the counts of modified and unmodified bases. **B.** m5C modification (<800 nt) landscapes. **C.** Ψ modification (>3000 nt) landscapes. **D.** Ψ modification (<800 nt) landscapes. With selection criteria based on single-base analysis (those that meet all the requirements are highlighted in vertical lines): For the positive strand with read length > 3000 nt, 0h vs. 20 minutes decrease, 20 minutes vs. 40 minutes increase (vertical blue line); 0h vs. 20 minutes decrease, 20 minutes vs. 40 minutes no difference (vertical purple line); 0h vs. 20 minutes increase, 20 minutes vs. 40 minutes no difference (vertical brown line); For the positive strand with read length <800 nt. 20 minutes vs. 40 minutes increase (vertical cyan line), 20 minutes vs. 40 minutes no difference (vertical red line). Stoichiometry (test samples) - stoichiometry (IVT samples) > 0, balanced MAP-based *p*-value < 0.05. All *p*-values pass the significance *q* value for reducing the False Discovery Rate (FDR). 5-methylcytidine, m^5^C.

We next assessed the secondary structures at these differentially methylated sites and the potential motifs that have served as substrates for known modification enzymes. We found no specific modification sites or secondary structures for m5C modification. We observed conserved loops or at the stem-loop junctions Ψ installation motif on the MS2 RNA (**Suppl Figure S10 A-L**). MEME motif pattern showed the conserved sequence among these sites (U-rich hairpin) (**Suppl Figure S10 M-N**). On a conserved position at 924 nt (with the highest stoichiometry: >80% among all the modification sites), we found a conserved recognition structure (installation at stem-loop junctions) and a highly conserved sequence compatible with ΨUACA RluA canonical recognition motif (Schaening-Burgos et al., 2024). Other pseudouridine installation sites exhibited relatively low stoichiometry (<30%) compared to location 924 (>80%). Analysis of their secondary structures showed either a fold compatible with ΨUACA RluA canonical recognition motif or a non-canonical motif substrate (**Suppl Figure. 10)**. Together, these results suggest that RNA modification was present in the initial transcripts and throughout the life cycle without the acquisition of new modified sites on full transcripts, whereas small transcripts do acquire and remove modified sites during replication. These newly modified sites, amenable to modifications, are U-rich hairpins for Ψ modification.

## Discussion

RNA bacteriophages such as Qβ and MS2 have been extensively studied for their ability to rapidly replicate (Chang et al., 2022; Vasilyev et al., 2013; Wagner, Weise, and Mutschler, 2022). However, the dynamics of their transcriptional activities across a single and multiple life cycles remain relatively unexplored. This study aimed to detail the dynamics of MS2 transcriptional activity using the Nanopore direct RNA sequencing approach across 6 hours post-inoculation. We found that MS2 phage maintained a high transcriptional activity, especially expressing high amounts of small transcripts containing the coat protein-encoding region. We also found hybrid transcripts and foldback RNA that have arisen from template switching recombination. By investigating the RNA modification landscapes, we found a conserved U-rich hairpin compatible with RluA-mediated Ψ modification. Together, direct RNA Nanopore sequencing has allowed the dissection of the detailed dynamics of MS2 transcriptional activity across time. This has led to our main finding that, despite carrying a minimal genome, MS2 efficiently enhanced its transcriptional activity and harnessed its error-prone replicase to rapidly attach, replicate, and lyse the bacterial cell.

### High transcriptional/replication level of MS2 within bacterial cells

Previous works assessed in depth RNA phages (Vasilyev et al., 2013) and RNA viruses (Hillen et al., 2020) replication patterns (including host factors). The archetypical MS2 and Qβ replicase complexes have been extensively investigated since the 1960s, including the host factors responsible for forming the complex (Wagner et al., 2022) and their underlying roles (Vasilyev et al., 2013). However, little is known about the dynamics of the transcriptional activity during the MS2 phage life cycles. A key reason is that most previous studies were performed in an *in vitro* or a transcription-translation system for MS2 (Wagner et al., 2022) and Qβ (Vasilyev et al., 2013; Tomita, Ichihashi, and Yomo, 2015). Using Nanopore RNA DRS, we reported significant changes in MS2 transcript levels across multiple life cycles. Strikingly, we noticed an over 50-fold increase in full-length MS2 transcripts in only three hours post-inoculation and over 400-fold increase in coat protein expression, indicative of high activity of the phage for replication and assembly to maintain the infection. To a lesser extent, other RNA viruses, such as the Human Coronavirus 229E (HCoV-229E), produce their own transcripts (∼33% reads mapped to the virus genome) at high levels (Viehweger et al., 2019) for viral replication. Another study also showed a large proportion of reads (∼65% reads) mapped to the severe acute respiratory syndrome coronavirus 2 (SARS-CoV-2) (Kim et al., 2020). As such, our observation is in line with previous reports on other viruses, suggesting the ability of a virus to rapidly transcribe within hours post-inoculation. Together, our study reports that RNA phages such as MS2 require significant upregulation and fine-tuning of the expression of their ORFs across time.

### Formation of subgenomic transcripts during the MS2 life cycle

Previous works have reported that MS2 utilizes its negative strand for replication via the replicase complex (Dykeman, 2023; Wagner, Weise & Mutschler, 2022). While the Qβ replicase complex has been extensively studied for the formation of spontaneous short amplifying RNA and additional by-products, MS2 replicase has been recently described as less promiscuous towards the formation of spontaneous replicative products (Wagner, Weise & Mutschler, 2022).

During our investigation, we identified a large population of short subgenomic reads appearing after the latent period of phage infection and increasing over time. We hypothesised that this population could be a Nanopore sequencing bias due to degraded RNA from bacterial lysis after MS2 infection, and 3’ end enrichment is the nanopore DRS 3’ end bias (Jain et al., 2022). We also hypothesised that these short transcripts could be single ORF transcripts. We found indeed evidence of RNA degradation and Nanopore DRS sequencing bias. Interestingly, we identified hotspots of mapped transcripts corresponding to the full-length coat protein ORF. Specifically, the top RNA virion contact sites were previously identified at positions surrounding 1740-1750 nt and 2110-2120 nt (Rolfsson et al., 2016) and overlayed with the top termination hits **(Figure 3D),** suggesting the assembly and packaging process could potentially inhibit the replication process by prematurely ending the transcription at specific RNA contact sites (**Figure 3E-F**). Additionally, we reported that this transcription is potentially regulated by the occupancy of the coat protein on the positive strand on specific RNA binding sites, potentially aborting the transcription/translation from the replicase complex (Rolfsson et al., 2016). Together, we suggest that MS2 phage produces a high level of coat protein transcripts to potentially repress the replicase complex, and critical for virion assembly.

### Error-prone replicase complex generates hybrid transcripts

An important requirement for some viruses to replicate and evolve is to harbour either proofreading (Robson et al., 2020) or non-proofreading replicase (Whitfield et al., 2020). A mechanism for viruses to acquire novel traits and evolve is to form hybrids with other viruses or within the same virus (Simon-Loriere and Holmes, 2011). We wondered whether the error-prone replicase would favour the generation of *de novo* mutations and such hybrid RNAs to form new virions. Using Nanopore DRS, we reported long intraspecies hybrid transcripts (>3569 nt), including fold-back RNAs accounted for 0.2% of the total transcript count. We also found that these hybrid transcripts resulted from template switching recombination. The presence of hybrid RNA or recombination events has been reported for other eukaryotic viruses such as SARS-CoV-2 (Garushyants, Rogozin, and Koonin, 2021) or plant viruses (Johnson et al., 2024). Hybrid RNAs have also been reported for MS2 in plasmid systems (van Meerten et al., 2002). Consistent with previous observations, our findings indicate the ability of MS2 to recombine using template switching mechanisms into new RNA transcripts.

### U-rich hairpin of Ψ installation sites on native MS2 RNA

Previous works have evidenced the crucial role of RNA modification in viral replication (Bhattacharya et al., 2022). To achieve high translation efficiency and evade host immunity, some viruses encode their own methyltransferase or other modifying enzymes (Li and Rana, 2022). However, the MS2 genome doesn’t harbor any modifying enzymes and requires hijacking the host to modify its own RNA. Using Nanopore DRS, we aimed to assess whether the modification landscape within a single life cycle and across time. We evidenced the presence of the Ψ installation site and secondary structure (U-rich hairpin) amenable to Ψ modification with a variation of stoichiometry within a single cycle and across time. We notably found evidence of canonical and non-canonical ΨUACA RluA-mediated installation, providing evidence that host pseudouridine synthase modifies MS2 RNA during its life cycle. Interestingly, we found a dynamic of loss and gain in methylation of the small transcripts during the MS2 across time. It suggests that, unlike the full transcripts, small transcripts are more amenable to RNA modification, indicating the distinct secondary structure of small transcripts compared to full-length transcripts, rendering them more amenable to Ψ installation. Together, it confirms previous work that the MS2 transcript is amenable to RNA modification (Jones, 2023). These modifications are either transient, predominantly on short transcripts, or permanent with various stoichiometries on canonical ΨUACA RluA-mediated installation sites. However, the functionality of these modifications on MS2 ORF translation remains unknown and will require further investigations.

## Conclusion

Our investigation, capturing the dynamics of MS2 transcriptional activity across time using the Nanopore direct RNA sequencing approach, has revealed a complex interplay between the replicase, the coat protein, and the negative RNA strand to control and fine-tune its transcription. It has revealed the numerous generations of byproducts during transcription, as either products of the error-prone replicase (hybrid reads) or coat protein transcripts required for assembly. We finally reported the dynamics of modified RNA stoichiometry changes during a single cycle and the addition of new modification sites with low stoichiometry of the canonical and non-canonical substrate motif RulA-mediated. Together, the direct RNA sequencing approach has revealed new insights into the complex transcriptional dynamics of MS2 bacteriophage.

## Acknowledgements

The authors would like to acknowledge the Biomolecular Resource Facility (Dr Carolina Correa-Ospina and Mr Ziyan Zhang) for assistance in the Nanopore sequencing. The authors would like to acknowledge Mr Agin Ravindran for helping with the magnetic beads purification. The authors would like to acknowledge the Burgio laboratory for comments on the early version of the manuscript. This work was undertaken with the assistance of resources from the National Computational Infrastructure (ANUMAS and NCMAS schemes), an NCRIS enabled capability supported by the Australian Government. GB work is supported from the National Health and Medical Research Council, the Medical Research Future Funds and the Australian Research Council and the Medical Research Future Fund.

## Authors contribution

GB conceived the study. GB supervised the work and provided funding. NN performed the experiments. NN and GB performed the analysis. NN and GB wrote the paper.

## Supplementary figures

**Supplementary Fig. S1.**
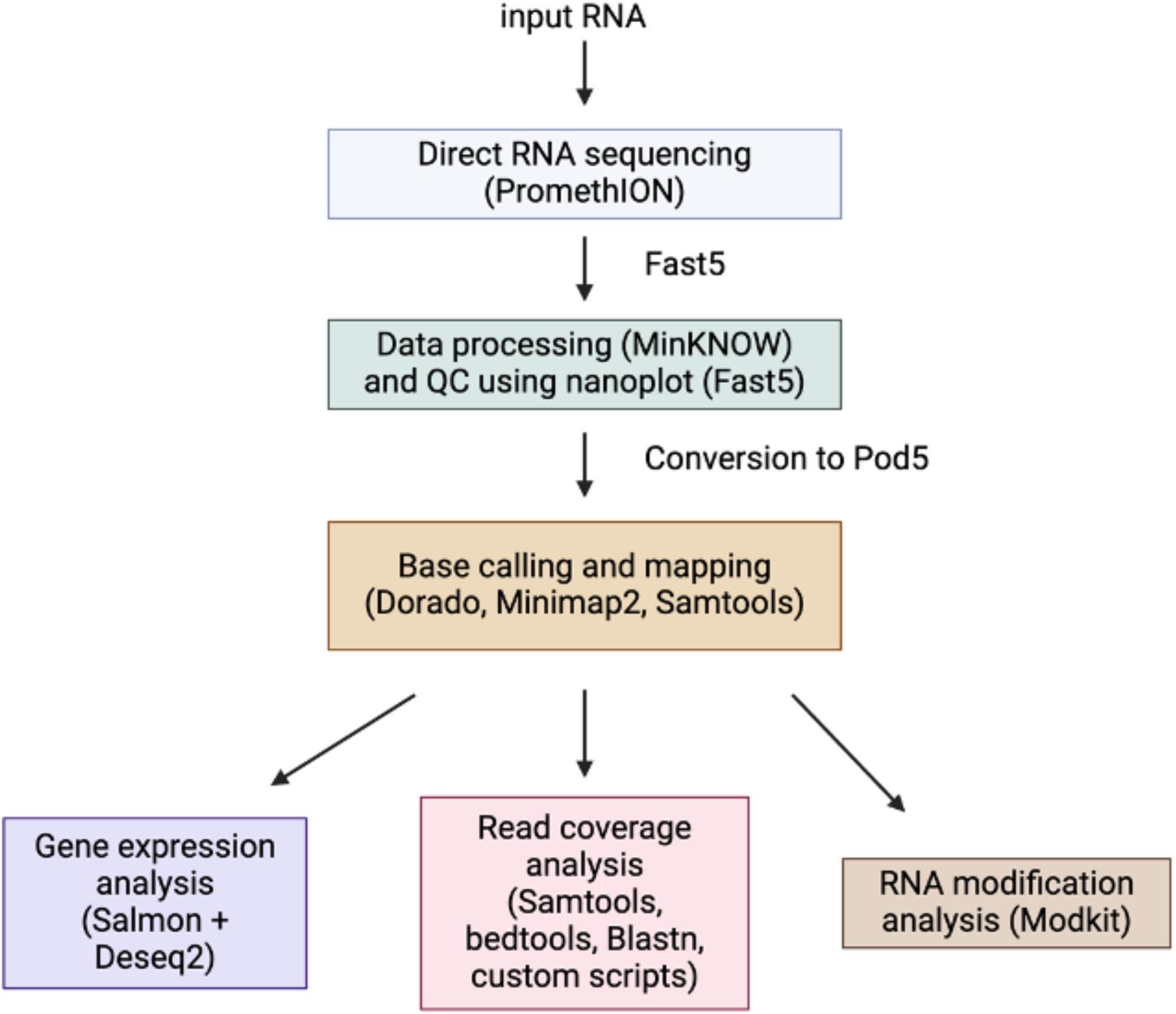
**The schematic computational analysis workflow for processing the Oxford RNA-direct Nanopore sequencing datasets for MS2 RNAs.**

**Supplementary Fig. S2.**
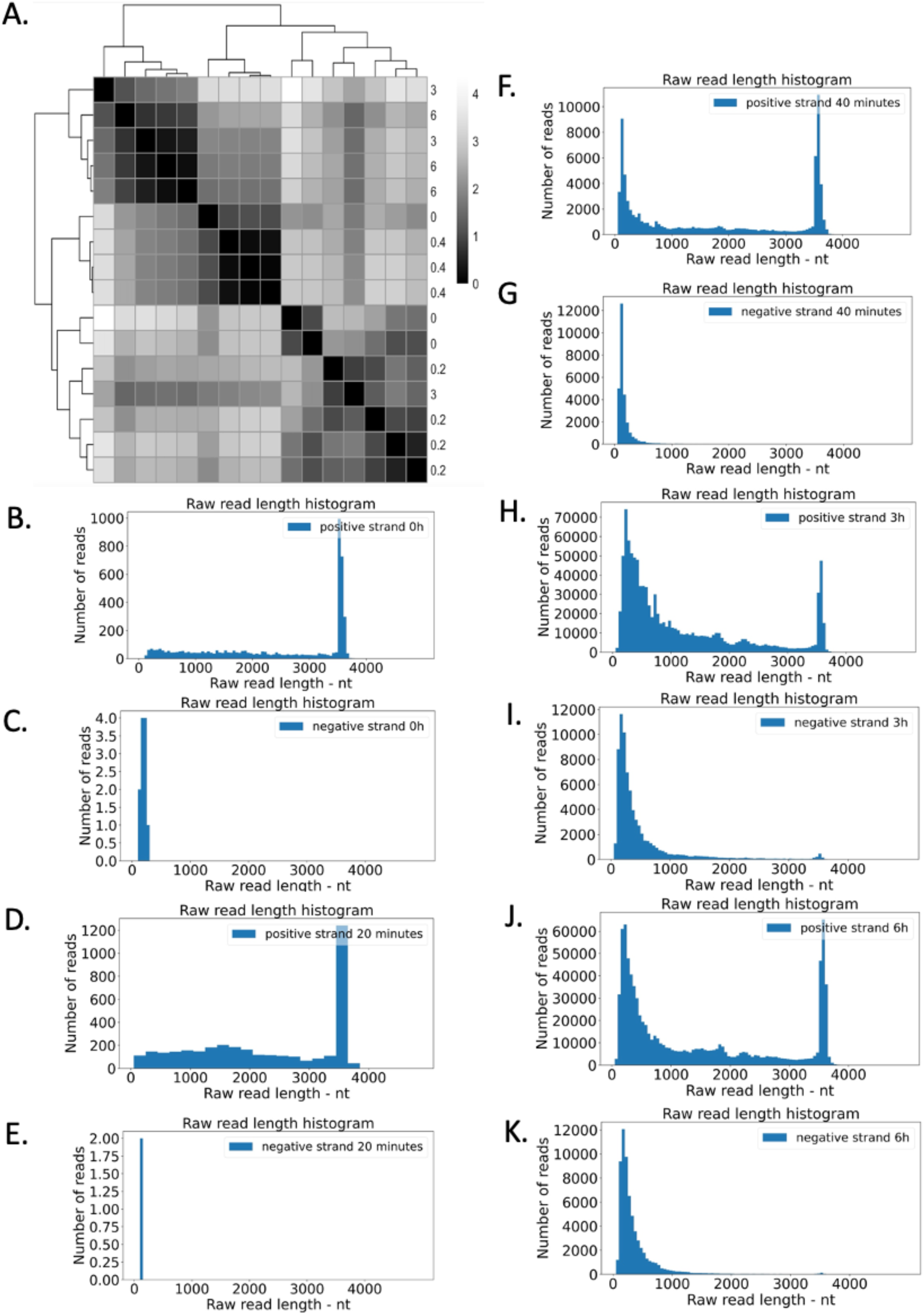
**A.** Heatmap of MS2 sample-to-sample distances. **B-I.** Raw read length distribution histogram of sequencing basecalled reads of MS2 samples **B.** positive strand at 0 h **C**. negative strand at 0h, **D**. positive strand at 20 mins **E**. negative strand at 20 mins **F**. positive strand at 40 mins **G**. negative strand at 40 mins. **H**. positive strand at 3h **I**. negative strand at 3h **J**. positive strand at 6h **K**. negative strand at 6h.

**Supplementary Fig. S3.**
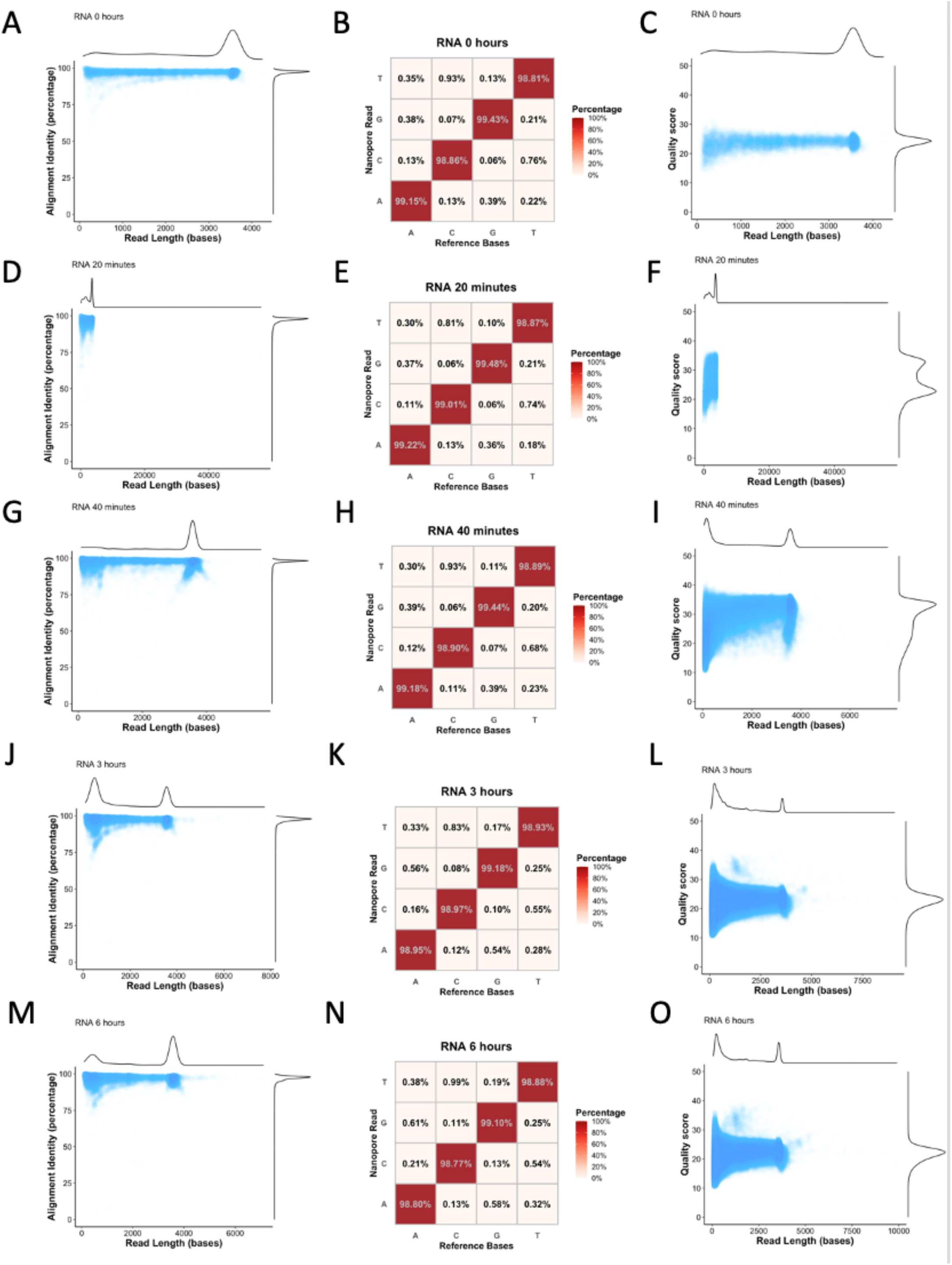
Performance metrics for Nanopore native RNA sequencing. Nanopore phage native RNA read alignment identity versus read length at different time points (0h, 20 minutes, 40 minutes, 3h, and 6h) (**Left**). Native RNA read base substitution matrix (**Middle**). The MS2 transcriptome’s known base identity is represented on the x-axis. For Nanopore reads, the y-axis is the base identity at the same position. Matches or mismatches are represented by the colour intensity in the boxes, with dark red standing for a high match of correct calls. Quality score versus read length at different time points (0h, 20 minutes, 40 minutes, 3h, and 6h) (**Right**). **A-C**. At 0h p.i. **D-F**. At 20 minutes p.i. **G-I**. At 40 minutes p.i. **J-L**. At 3h p.i. **M-O**. At 6h p.i.

**Supplementary Fig. S4.**
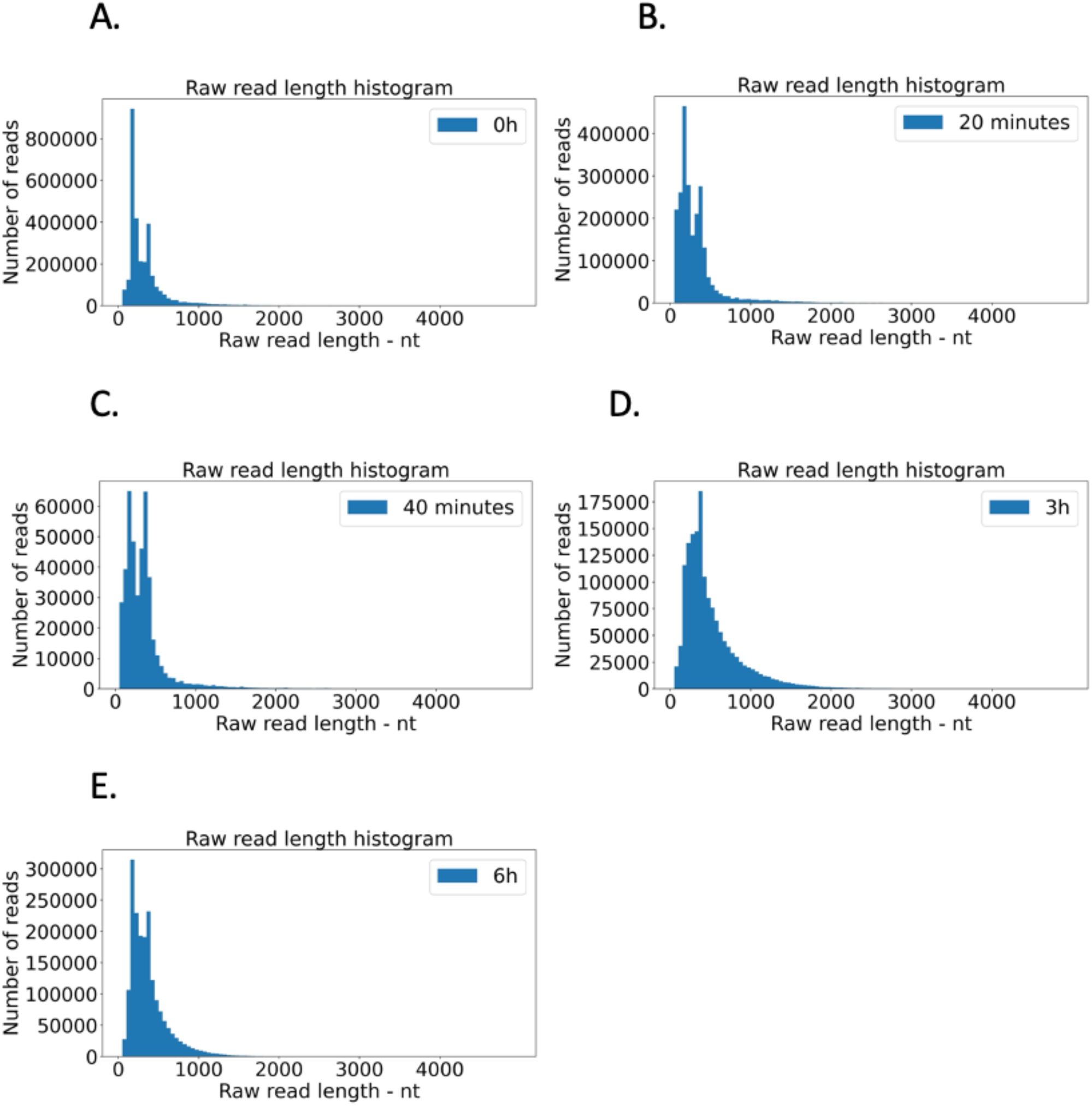
Raw read length distribution histogram of sequencing basecalled reads of host bacterial RNA (**A**) at 0h p.i., (**B**) 20 mins p.i., (**C**) 40 mins p.i., (**D**) 3h p.i., (**E**) 6h p.i.

**Supplementary Fig. S5.**
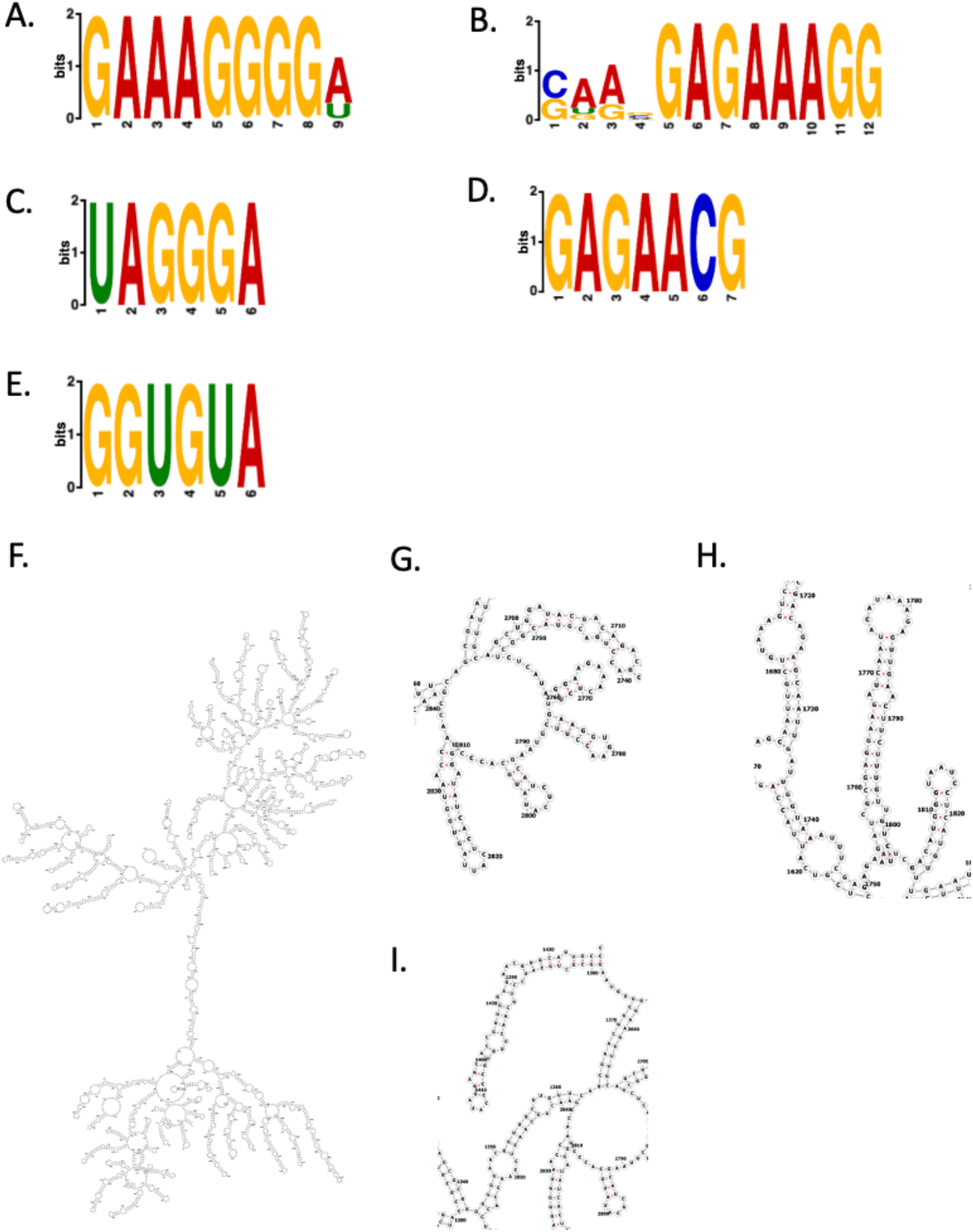
MEME alignment between putative replicase binding sites on MS2 negative strand and identified RBS sequence of genes in *Escherichia coli* K-12 MG1655 and secondary structure of template strand (negative strand) corresponding to the potential TES sites (positive-strand transcript). **A.** 8-16 nt, **B**. 1003-1010 nt, **C**. 1140-1145 nt, **D**. 1356-1364 nt, **E**. 1509-1514 nt are corresponding genomic locations on MS2 positive-strand, the conserved sequence showed the negative strand sequence from 5’ end to 3’ end on those locations. **F.** Secondary structure of native MS2 full-length negative strand, generated by RNAfold. The secondary structure showed the location of the negative strand at **G**. 2740-2840 nt, **H**. 1720-1820 nt, **I**. 1335-1435 nt, corresponding to the read end site, generated by RNAfold.

**Supplementary Fig. S6.**
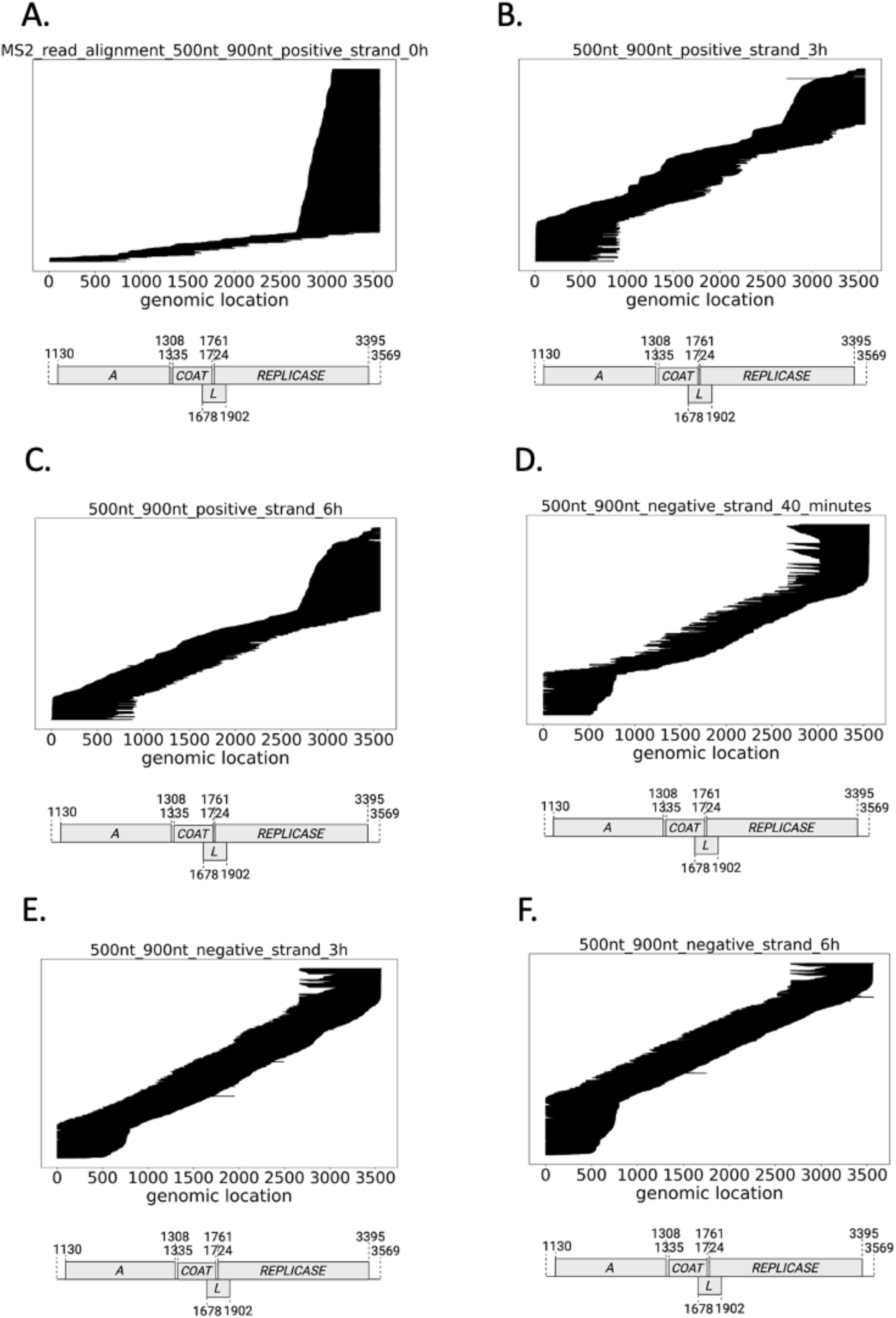
Reads aligned to MS2 genome (shown in lines), positive-strand or negative-strand, separated into different fractions of similar read length (500-900 nt) at 0h, 20 minutes, 40 minutes, 3h, and 6h p.i. **A.** Read aligned to positive-strand at 0h p.i., **B.** Read aligned to positive-strand at 3h p.i., **C.** Read aligned to positive-strand at 6h p.i., **D.** Read aligned to negative-strand at 40 minutes p.i., **E.** Read aligned to negative-strand at 3h p.i., **F.** Read aligned to negative-strand at 6h p.i.

**Supplementary Figure S7.**
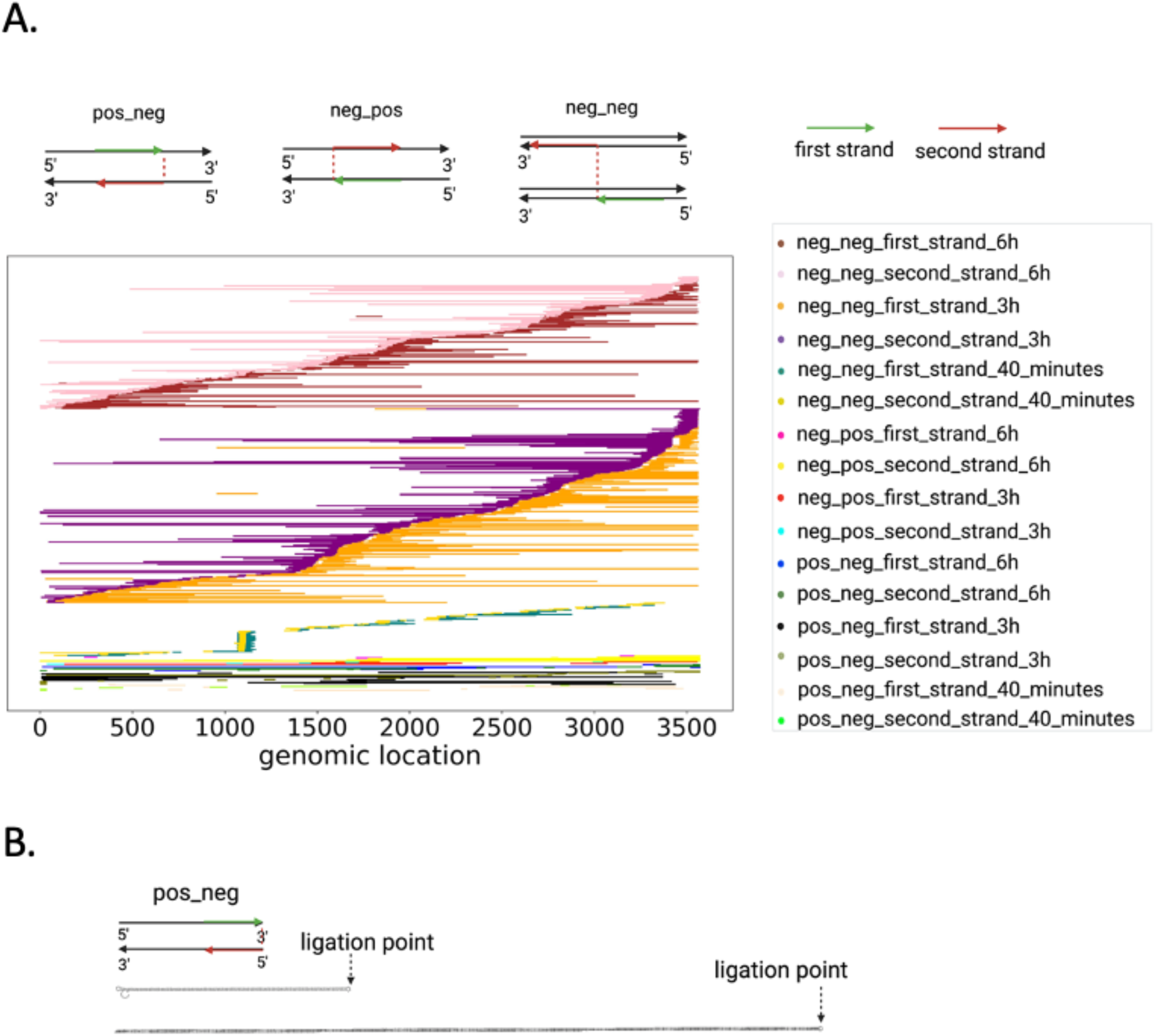
Systematically summarised the characteristics of MS2 hybrid reads. **A.** Features of MS2 hybrid reads of neg_neg, pos_neg, and neg_pos. Total hybrid reads were visualised. **B.** Schematic diagram and secondary structure of fold-back RNAs (3’ end of the positive strand ligates with 5’ end of the negative strand start) by RNAfold. Positive-strand 3451-3568 ligated with negative-strand 2-135 (positive-strand location 3568-3435), and positive-strand 3173-3568 ligated with negative-strand 2-395 (positive-strand location 3568-3175).

**Supplementary figure S8.**
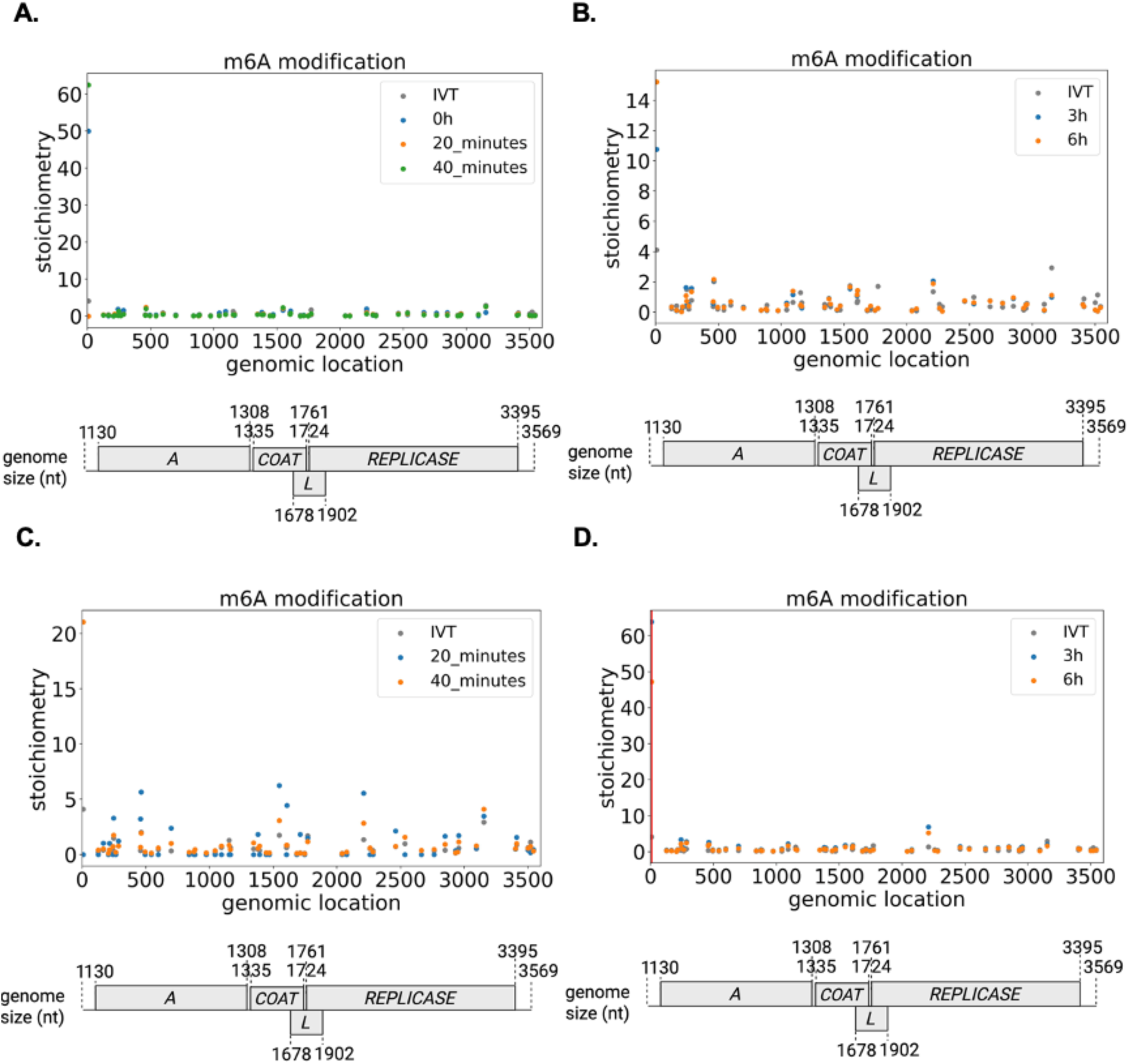
Frequent potential sites of m^6^A (based on DRACH motifs) RNA modification dynamics on MS2 phage positive RNA (>3000 nt transcript length and <800 nt transcript length) via Dorado and Modkit software packages. Synthetic modification-free RNA (IVT RNA) as the negative control. **A.** m6A (DRACH) modifications (>3000 nt) at 0h, 20 minutes, and 40 minutes **B.** m6A (DRACH) modifications (>3000 nt) at 3h and 6h. **C.** m6A (DRACH) modifications (<800 nt) at 0h, 20 minutes, and 40 minutes **D.** m6A (DRACH) modifications (<800 nt) at 3h and 6h. With selection criteria based on single-base analysis (those that meet all the requirements are highlighted in vertical lines): Presence of modification at 3h and 6h (vertical red line). stoichiometry (test samples) - stoichiometry (IVT samples) > 0, balanced MAP-based *p*-value < 0.05. All *p*-values pass the significance *q* value for reducing the False Discovery Rate (FDR).

**Supplementary figure S9.**
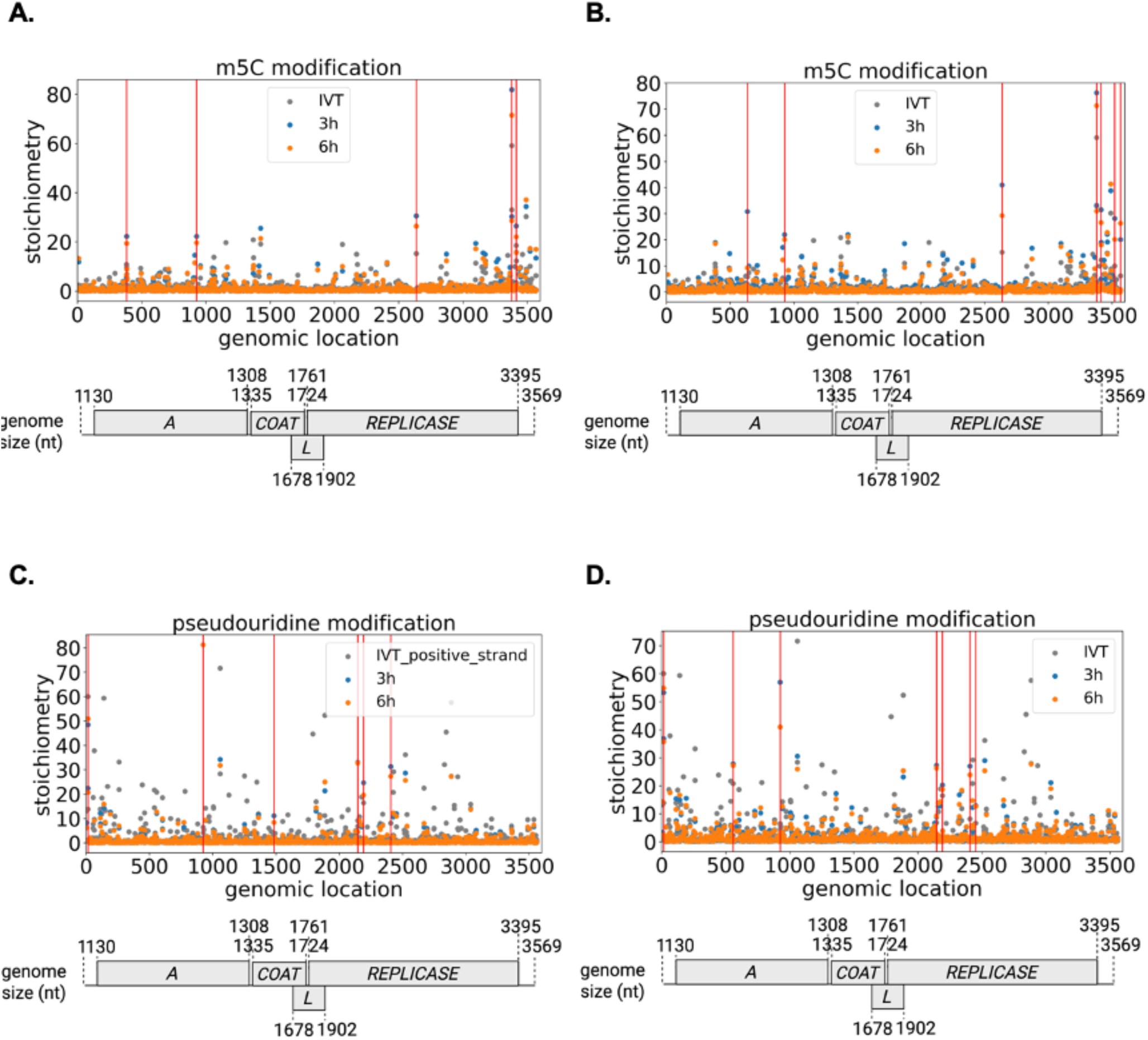
Frequent potential sites of Ψ and m^5^C RNA modification dynamics on MS2 phage positive RNA (>3000 nt transcript length and <800 nt transcript length) via Dorado and Modkit software packages. Synthetic modification-free RNA (IVT RNA) as the negative control. **A.** m5C modification (>3000 nt) landscapes. **B.** m5C modification (<800 nt) landscapes **C.** Ψ modification (>3000 nt) landscapes **D**. Ψ modification (<800 nt) landscapes. With selection criteria based on single-base analysis (those that meet all the requirements are highlighted in vertical lines): presence of modifications at 3h or 6h (vertical red line). Stoichiometry (test samples) - stoichiometry (IVT samples) > 0, balanced MAP-based *p*-value < 0.05. All *p*-values pass the significance *q* value for reducing the False Discovery Rate (FDR).

**Supplementary Figure S10.**
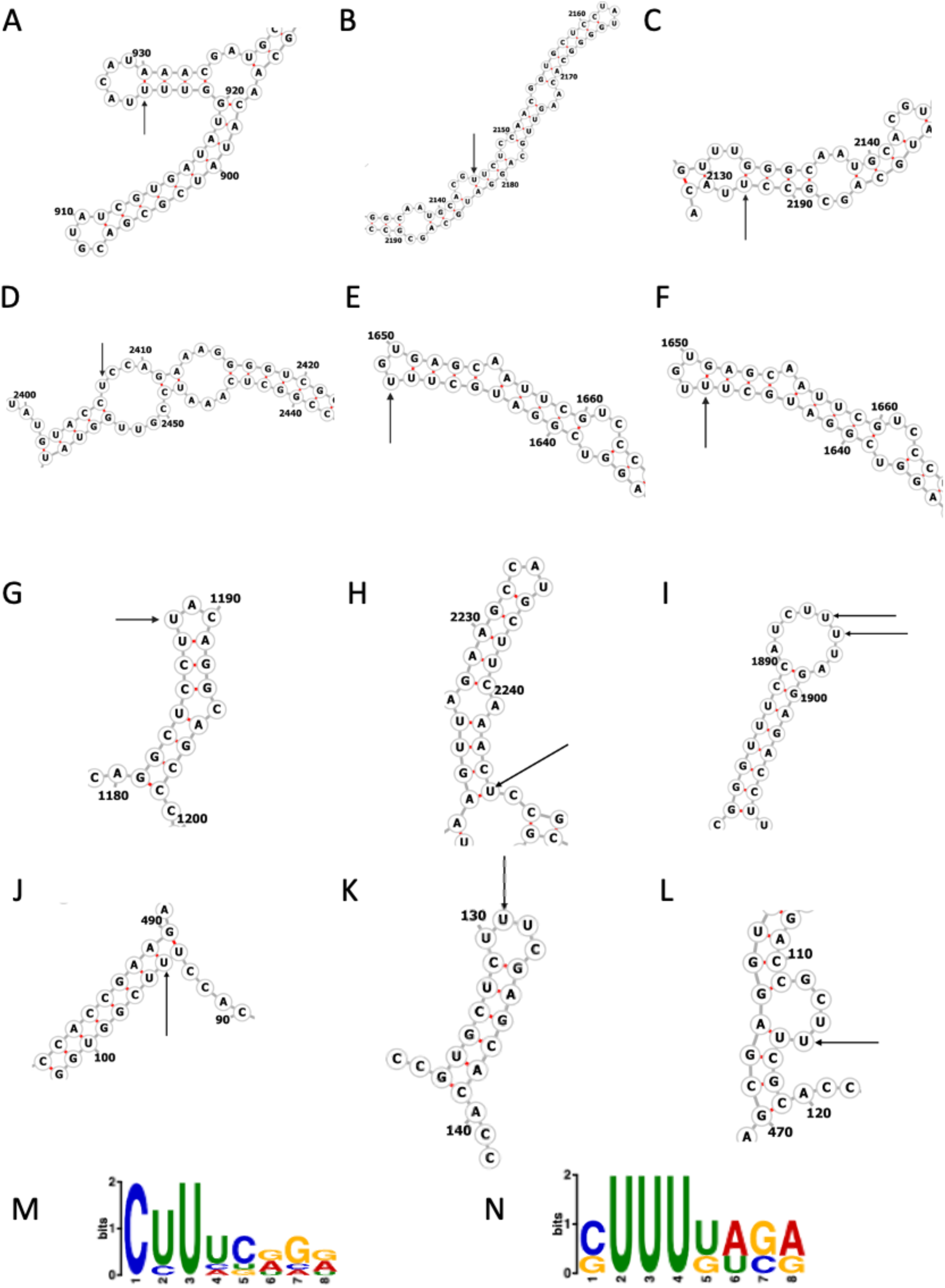
Conserved secondary structure of the installation of potential Ψ **modifications.** Positive strand. **A**. at location 924, **B**. at 2145, **C**. at 2192, **D**. at 2407. Negative strand at location **E**. 1922 (1648), **F.** 1923 (1647), **G**. 2382 (1188), **H**. 1325 (2245), **I**. 1674 (1896), 1675 (1895), **J**. 3475 (95), **K**. 3439 (131), **L**. 3455 (115). **M.** Conserved recognition sequence for D, F, G, I, J, K, and L, shown in MEME motif, position 3 is Ψ the site. **N**. Conserved recognition sequence for A, E, I, position 4 is Ψ the site. Position 1, 5’ end, position 8, 3’ end.

## Supplementary Tables

**Table S1:**
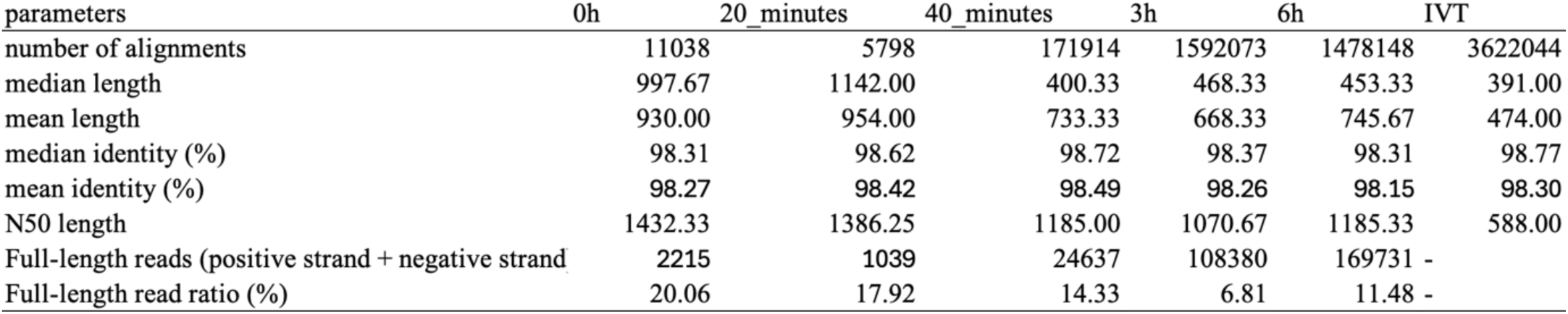
Oxford Nanopore Technologies (ONT) DRS statistical metric for read quality control.

| parameters | 0h | 20_minutes | 40_minutes | 3h | 6h | IVT |
| --- | --- | --- | --- | --- | --- | --- |
| number of alignments | 11038 | 5798 | 171914 | 1592073 | 1478148 | 3622044 |
| median length | 997.67 | 1142.00 | 400.33 | 468.33 | 453.33 | 391.00 |
| mean length | 930.00 | 954.00 | 733.33 | 668.33 | 745.67 | 474.00 |
| median identity (%) | 98.31 | 98.62 | 98.72 | 98.37 | 98.31 | 98.77 |
| mean identity (%) | 98.27 | 98.42 | 98.49 | 98.26 | 98.15 | 98.30 |
| N50 length | 1432.33 | 1386.25 | 1185.00 | 1070.67 | 1185.33 | 588.00 |
| Full-length reads (positive strand + negative strand) | 2215 | 1039 | 24637 | 108380 | 169731 | - |
| Full-length read ratio (%) | 20.06 | 17.92 | 14.33 | 6.81 | 11.48 | - |

**Table S2:** Read counts from Nanopore direct RNA sequencing of total RNA from DSM5S95 *E coli* cells infected with MS2 phage at different time points.

Number of alignments
| Mapped to | MS2 | Bacterial genome | Bacterial plasmid | Total | MS2/Total*100% | Bacterial genome/Total*100% | Bacterial plasmid/Total*100% | MS2/Bacterial genome |
| --- | --- | --- | --- | --- | --- | --- | --- | --- |
| 0h | 33114 | 1939187 | 55796 | 2028097 | 1.63% | 95.62% | 2.75% | 1.71% |
| 20 minutes | 21414 | 1938841 | 83573 | 2043828 | 1.05% | 94.86% | 4.09% | 1.10% |
| 40 minutes | 515742 | 390761 | 12380 | 918883 | 56.13% | 42.53% | 1.35% | 131.98% |
| 3h | 4776219 | 1547128 | 52017 | 6375364 | 74.92% | 24.27% | 0.82% | 308.72% |
| 6h | 4434444 | 1738813 | 46870 | 6220127 | 71.29% | 27.95% | 0.75% | 255.03% |

**Table S3:**
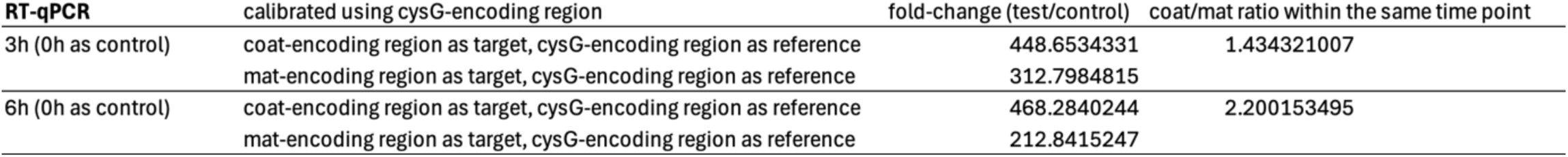
RT-qPCR of MS2 phage genes at different time points.

**Table S4:**
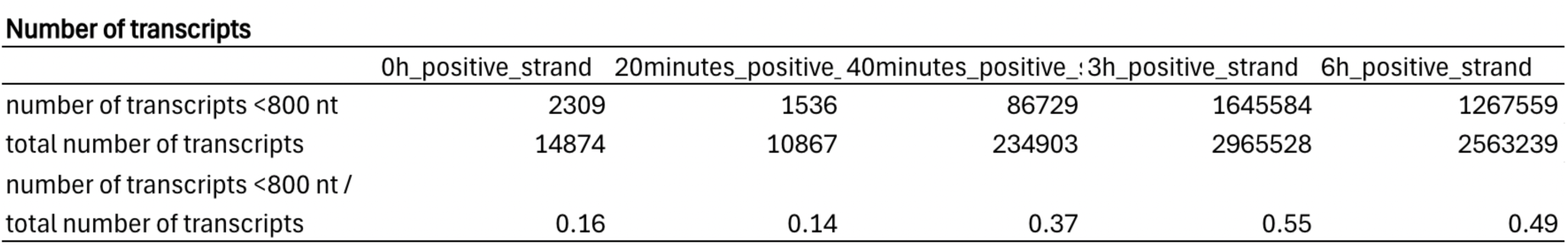
Number of transcripts (positive strand) with read length <800 nt.

**Table S5:** Number of transcripts with read length ranging from 100 nt to 800 nt.

Number of transcripts
|  | 0h | 20 mins | 40 mins | 3h | 6h |
| --- | --- | --- | --- | --- | --- |
| Number of transcripts with length ranging 100 nt -800 nt | 2765093 | 1979368 | 370390 | 1270874 | 1744140 |
| Number of total transcripts | 3031068 | 2352732 | 434632 | 1543057 | 1891854 |
| proportion (%) (Number of transcripts with length ranging 100 nt -800 nt/Number of total transcripts) | 91.23 | 84.13 | 85.22 | 82.36 | 92.19 |

**Table S6:**
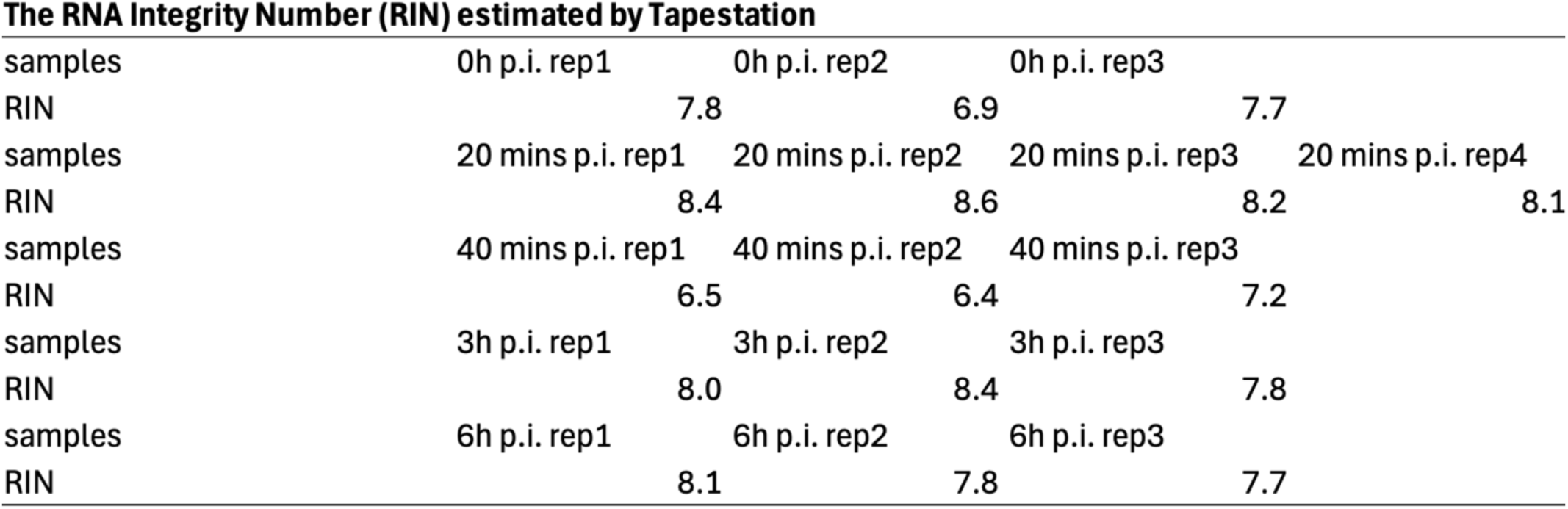
The RNA Integrity Number (RIN) per sample estimated by tapestation.

**Table S7:** Number of transcripts covering the coat protein-encoding region with read length <800 nt at the 3h time point.

Number of transcripts
|  | Reads (*coat* ORF) | Reads (total) | Percentage (%) |
| --- | --- | --- | --- |
| Transcripts from positive-strand with read length <800 nt at 3h p.i. | 620092480 | 116629530 | 18.81% |

**Table S8:**
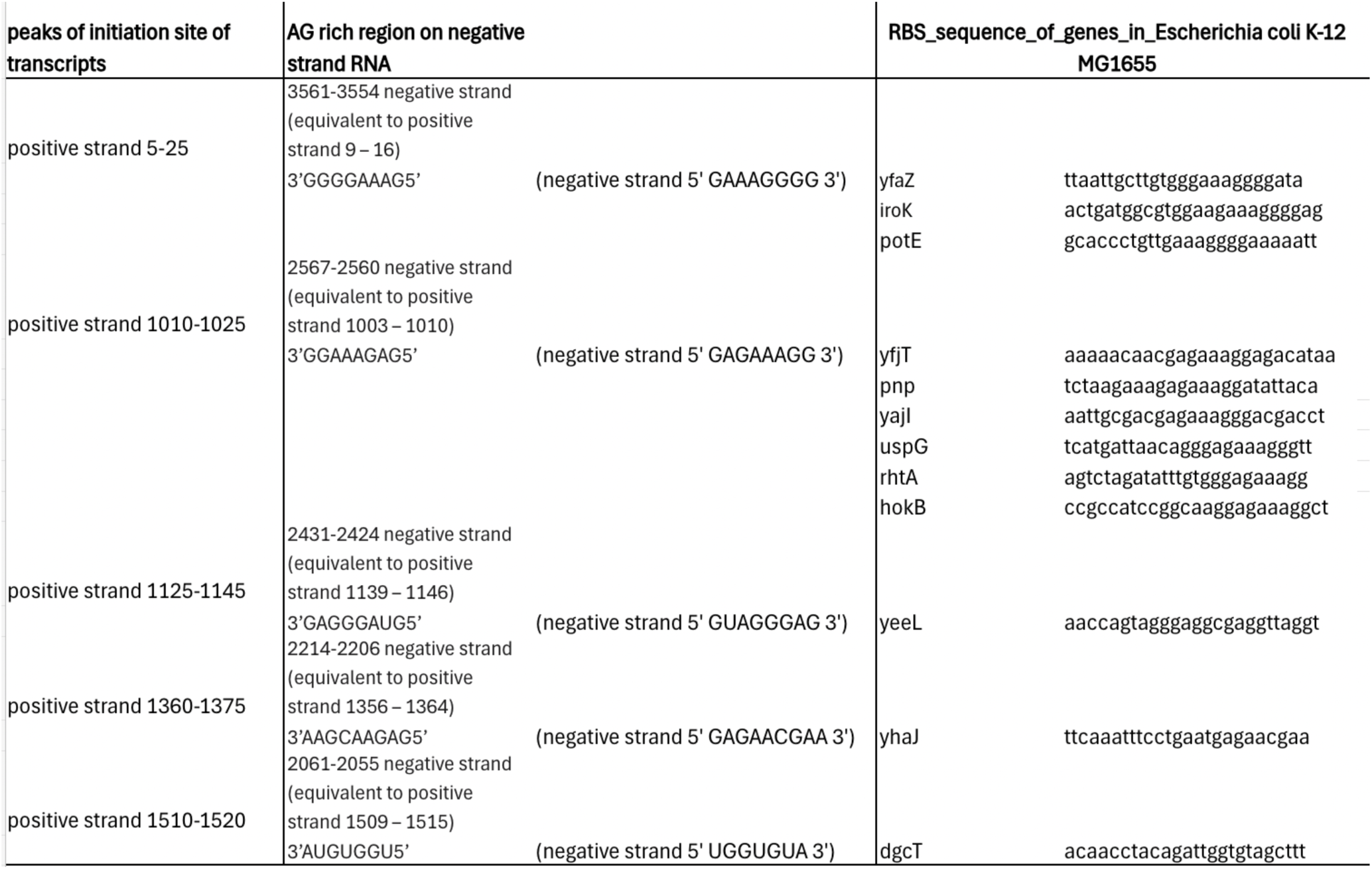
Putative replicase binding sites on MS2 negative strand and identified RBS sequence of genes in Escherichia coli K-12 MG1655.

**Table S9:** Number of transcripts with read length ranging from 500 nt to 900 nt aligned to different regions on the MS2 genome (positive strand) at different time points.

| read length ranging from 500 nt to 900 nt (For positive strand) |  |  |  |  |  |
| --- | --- | --- | --- | --- | --- |
|  | 0h | 20 mins | 40 mins | 3h | 6h |
| number of alignment (total) | 1040 | 732 | 15107 | 15901 | 20471 |
| number of alignment (mapped to the 3' end)(enrichment due to Nanopore sequencing 3' bias) | 866 | 574 | 6424 | 3172 | 5996 |
| proportion (%) (number of alignment (mapped to the 3' end)/number of alignment (total)*100%) | 83.27 | 78.42 | 42.52 | 19.95 | 29.29 |
| number of alignment (mapped to the 5' end)(degraded RNA) | 21 | 39 | 4583 | 3075 | 2290 |
| proportion (%) (number of alignment (mapped to the 5' end)/number of alignment (total)*100%) | 2.02 | 5.33 | 30.34 | 19.34 | 11.19 |
| number of alignment (mapped to the coat protein encoding region) | 20 | 20 | 511 | 389 | 842 |
| proportion (%) (number of alignment (mapped to the coat protein encoding region)/number of alignment (total)*100%) | 1.92 | 2.73 | 3.38 | 2.45 | 4.11 |
| number of alignment (mapped to the lysis protein encoding region) | 22 | 11 | 828 | 1773 | 1630 |
| proportion (%) (number of alignment (mapped to the lysis protein encoding region)/number of alignment (total)*100%) | 2.12 | 1.50 | 5.48 | 11.15 | 7.96 |
| number of alignment (mapped to the coat and lysis protein encoding region) | 6 | 1 | 186 | 311 | 216 |
| proportion (%) (number of alignment (mapped to the coat and lysis protein encoding region)/number of alignment (total)*100%) | 0.58 | 0.14 | 1.23 | 1.96 | 1.06 |

**Table S10:** Number of transcripts with read length ranging from 500 nt to 900 nt aligned to different regions on the MS2 genome (negative strand) at different time points.

Read length ranging from 500 nt to 900 nt (For negative strand)
|  | 40 mins | 3h | 6h |
| --- | --- | --- | --- |
| number of alignment (total) | 1708 | 18074 | 23389 |
| number of alignment (mapped to the 5' end of negative strand)(degraded RNA) | 465 | 2851 | 2459 |
| proportion (%) (number of alignment (mapped to the 5' end)/number of alignment (total)*100%) | 27.22 | 15.77 | 10.51 |
| number of alignment (mapped to the 3' end of negative strand)(enrichment due to Nanopore sequencing 3' bias) | 156 | 1078 | 2362 |
| proportion (%) (number of alignment (mapped to the 3' end)/number of alignment (total)*100%) | 9.13 | 5.96 | 10.10 |

**Table S11:** Number of deletion and duplication events.

Number of hybrid reads
|  | 20 minutes | 40 minutes | 3h | 6h |
| --- | --- | --- | --- | --- |
| deletions | 22 | 68 | 3335 | 3848 |
| duplications | 42 | 262 | 5417 | 5585 |

**Table S12:**
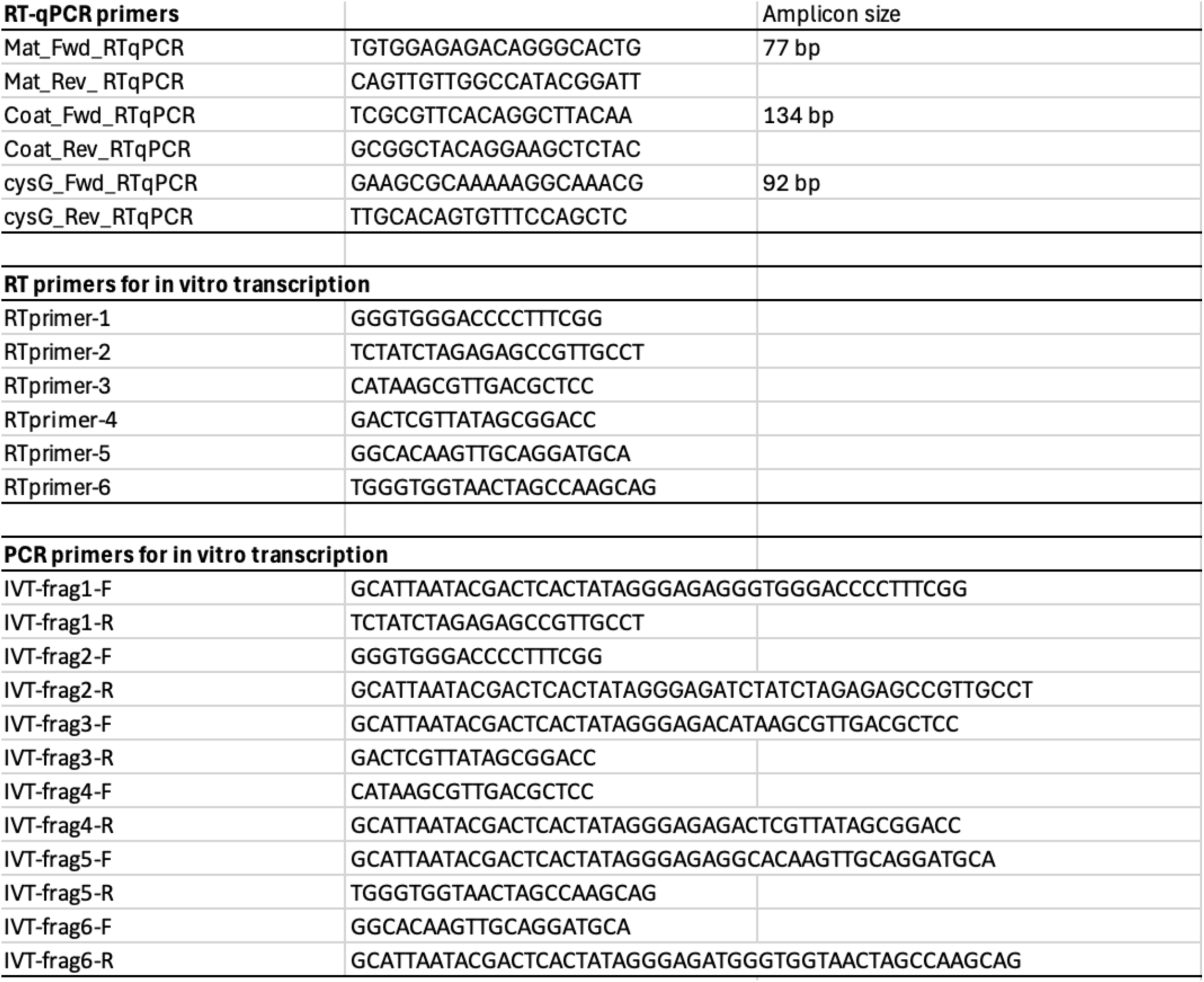
Primers used in the study.

